# Alcohol Withdrawal Alters the Inhibitory Landscape of the Prelimbic Cortex in an Interneuron- and Sex-specific Manner

**DOI:** 10.1101/2024.11.19.624401

**Authors:** Sema G Quadir, S. Danyal Zaidi, Meredith G Cone, Sachin Patel

**Author notes:** **Corresponding Author**: Sema G Quadir.

## Abstract

Alcohol use disorder (AUD) is highly prevalent and associated with substantial morbidity and high mortality among substance use disorders. While there are currently three FDA-approved medications for treating AUDs, none specifically target the withdrawal/negative affect stage of AUD, underscoring the need to understand the underlying neurobiology during this critical stage of the addiction cycle. One key region involved in alcohol withdrawal and negative affect is the prelimbic cortex, a subregion of the medial prefrontal cortex. While previous studies have examined alcohol-related adaptations in prefrontal cortical principal glutamatergic neurons, here we used male and female PV:Ai14, SOM:Ai14 and VIP:Ai14 mice to examine synaptic adaptations in all three major classes of prelimbic cortex interneurons following 72 hour withdrawal from a continuous access to two bottle choice model of EtOH drinking in male and female mice. We found that alcohol withdrawal increased excitability of prelimbic PV interneurons in males, but decreased excitability in prelimbic VIP interneurons in females. Additionally, alcohol withdrawal reduced GABA release onto PV interneurons in males while increasing glutamate release onto VIP interneurons in females. In SOM interneurons, alcohol withdrawal had no effect on excitability, but decreased glutamate release onto SOM interneurons in males. Together, our studies identified sex-specific alcohol withdrawal-induced synaptic plasticity in three different types of interneurons and could provide insight into the cellular substrates of negative affective states associated with alcohol withdrawal.

## INTRODUCTION

In 2023, 132.9 million adults in the US reported drinking alcohol (EtOH) in the last month with over 28 million adults meeting the criteria for an alcohol use disorder (AUD)^1^. AUD is traditionally conceptualized as a three-stage cycle –binge/intoxication, withdrawal/negative affect, and preoccupation/anticipation (see ^2–4^ for review). Chronic EtOH consumption shifts the motivation to consume EtOH from positive reinforcement (i.e. drinking to induce euphoria) to negative reinforcement (i.e. drinking to alleviate negative affect). Currently, three FDA-approved medications are available for AUD^5–8^; however, none specifically target the withdrawal/negative affect stage, emphasizing the need to understand neuroadaptations that occur during this stage of the addiction cycle.

Furthermore, despite the growing recognition of AUD’s impact on both men and women, most preclinical and clinical research has included predominantly male subjects. However, women progress more rapidly from casual drinking to AUD, meet more diagnostic criteria for the disorder, and experience quicker declines in health^9–17^. Historically, the rates of AUD have been higher in men than in women, but this gap is steadily narrowing. In 2013, the National Survey on Drug Use and Health reported 10.8 million men and 5.8 million women met the criteria for AUD ^18^. By 2023, those numbers increased to 14.6 million men and 10.6 million ^1^ women, resulting in a decrease of the proportion of men within the total AUD population from 65% to 58%. These statistics also demonstrate increases in total diagnoses over time, making it crucial to investigate the underlying neural mechanisms responsible for this disorder.

One important brain region involved in AUD pathophysiology is the medial prefrontal cortex (mPFC). Clinical studies comparing controls and individuals with AUD have identified differences in both resting state and cue-induced activity in the mPFC, with the degree of mPFC dysfunction correlating with poorer treatment outcomes^19–25^. Rodent studies have identified a plethora of EtOH-induced alterations in mPFC structure and function^26–52^, including changes in neuronal excitability and synaptic transmission. However, most of the rodent studies, particularly those with an electrophysiology component, focus on pyramidal neurons despite interneurons constituting 10-20% of cortical neurons^53^. Given the dynamic control interneurons exert over pyramidal neuron responses^54^, targeting interneurons could reveal novel therapeutic strategies for AUD^55^. While interneurons have been linked to EtOH responses in a variety of brain regions^56–65^, only more recently have investigators delved into the prelimbic subregion of the mPFC. For instance, a recent study found that extended abstinence from binge drinking (6+ months) increased glutamate release and synaptic drive onto undefined interneurons in the mPFC^29^.

Interneurons can be classified into three groups based on their expression of specific markers: parvalbumin (PV, ∼ 40% of cortical interneurons), somatostatin (SOM, ∼ 30%), or the 5HT3a receptor (5HT3AR, ∼ 40%)^66^. Among these, 5HT3AR neurons can further be divided into vasoactive intestinal peptide expressing (VIP, 40% of 5HT3AR neurons, 12% of all interneurons) and non-VIP expressing neurons.

PV interneurons are typically fast spiking, exhibit low input resistance, and show minimal spike frequency adaptation^54, 66, 67^. These interneurons synapse onto both the soma and proximal dendrite of pyramidal neurons and exert powerful feedforward inhibition ^66, 68^. While mPFC PV interneurons were classically studied in the context of stress and fear^69–75^, emerging evidence suggests both acute (24h) and protracted (3-9d) EtOH withdrawal increases both PV interneuron excitability and excitatory drive^51, 52^.

SOM interneurons are characterized by their lower resting membrane potential, high input resistance, low frequency spiking, and the presence of spike frequency adaptation ^54, 66, 67, 76^. SOM interneurons primarily synapse on dendritic tufts of pyramidal neurons, thus modulating synaptic integration of incoming signals and participating predominantly in feedback inhibition^66, 77^. In addition to mediating negative affect and working memory ^72, 74, 78, 79^, mPFC SOM neurons are less excitable and exhibit reduced synaptic drive after acute (24h) EtOH withdrawal ^33, 51^ .

VIP interneurons exhibit irregular spiking patterns, high input resistance, and presence of spike frequency adaptation ^66, 67^. Unlike PV and SOM interneurons, VIP interneurons predominantly inhibit interneurons rather than pyramidal neurons and thus act primarily via disinhibition. VIP interneurons mainly inhibit SOM neurons, but can also inhibit PV interneurons to some extent^80–82^. VIP interneurons are involved in cognition, pain, and negative affect^82–84^. Indeed, acute (24h) EtOH withdrawal reduced excitability of VIP interneurons^50^.

The studies presented here examined the effects of 72h EtOH withdrawal — a time point which we have previously shown to be associated with negative affective states and hyperalgesia in male and female mice^85^ — on the excitability and synaptic transmission of prelimbic cortex (PL) interneurons in , using male and female transgenic mice expressing fluorescent tdTomato in PV, SOM, or VIP interneurons.

## METHODS

### Animals

Male and female PV:Ai14, SOM:Ai14, and VIP:Ai14 mice were generated by crossing PV-IRES-Cre (JAX stock #017320), SOM-IRES-Cre (JAX stock #013044), and VIP-IRES-Cre (JAX stock #010908) mice with Ai14 (JAX stock #007914) mice, thus allowing for tdTomato expression in the targeted interneuron subtype. All mice were bred on a consistent C57BL/6J background. Sex was assigned based on the presence of external genitalia. Mice were housed in standard cages with cotton nesting material and a plastic toy for environmental enrichment, in a temperature-controlled room maintained on a 14-hour light/8-hour dark cycle (lights on at 6:00 am, off at 8:00 pm). Food (Teklad irradiated diet LM-485) and water were provided ad libitum. To avoid litter confounds, mice from at least two litters per genotype were used. All experimental procedures adhered to the National Institutes of Health Guidelines for the Care and Use of Laboratory Animals and were approved by Northwestern University’s Institutional Animal Care and Use Committee.

### Continuous Access to 2 Bottle Choice (CA2BC) Model of Ethanol Drinking

Adult mice were acclimated to single housing and two water bottles for 7 days prior to the initiation of drinking experiments. Male and female mice were run in separate cohorts, and thus the results are presented separately. Mice were at 10 weeks old on average when single-housing began and were randomly assigned into water or EtOH drinking groups. Following acclimation to single housing, one water bottle was replaced with gradually increasing concentrations of EtOH, as described previously ^85, 86^. Briefly, mice were allowed to drink 3% (*v/v*) EtOH for 4 days, followed by 7% (*v/v*) EtOH for 7 days, and then 10% (*v/v*) EtOH for at least 21 days. EtOH solutions were prepared by diluting 95% EtOH in tap water: a 3% solution was made by mixing 30 mL of 95% EtOH with 970 mL of water; a 7% solution was created by combining 70 mL of 95% EtOH with 930 mL of water; and a 10% solution was prepared by adding 100 mL of 95% EtOH to 900 mL of water.

Every week, both bottle weights and mouse body weights were recorded, and the position of the EtOH bottle was alternated to control for side preference. This weighing occurred between 9:00 and 10:00 am, corresponding to 2-3 hours into the light cycle. To account for spillage, the weights of EtOH and water bottles from an empty cage (without an animal) were measured, and any spilled liquid was subtracted from the weekly intake total. Following spillage correction, EtOH preference was calculated as [mL EtOH consumed/(mL EtOH consumed + mL water consumed]*100%]. To examine the effects of EtOH withdrawal, EtOH bottles were removed and replaced with an additional water bottle 72h prior to being sacrificed for electrophysiology studies. This 72h timepoint was chosen based on our previous studies, which showed that withdrawal symptoms like hyperalgesia and irritability emerge at this specific timepoint following CA2BC^85^.

Control water-drinking mice, which were also single-housed for a similar duration, had continuous access to two water bottles throughout the experiment. These mice were weighed weekly, and their water bottles were handled in the same manner to ensure that both groups experienced equivalent levels of handling and cage disruption.

### Electrophysiology

Electrophysiology experiments were conducted as previously before^87, 88^. Briefly, mice were anesthetized with isoflurane and subsequently underwent transcardial perfusion with ice-cold, oxygenated (95% O₂, 5% CO₂) N-methyl-D-glucamine (NMDG)-based artificial cerebrospinal fluid (ACSF)^89^. The composition of this ACSF included (in mM): 93 NMDG, 2.5 KCl, 1.2 NaH₂PO₄, 30 NaHCO₃, 20 N-2-hydroxyethylpiperazine-N-2-ethane sulfonic acid (HEPES), 25 glucose, 5 sodium ascorbate, 3 sodium pyruvate, 5 N-acetylcysteine, 0.5 CaCl₂·4H₂O, and 10 MgSO₄·7H₂O (pH 7.30-7.32, osmolarity 297-298 mOsm). Following the perfusion, the brain was swiftly removed, and 250-μm coronal slices containing the PL were prepared using a vibratome (Leica Biosystems, model # VT1000S) while immersed in the NMDG solution. These slices were incubated in oxygenated NMDG-ACSF at 32 °C for 10-15 minutes before being stored at 24 °C until recordings were performed in HEPES-based ACSF, which was comprised of (in mM): 92 NaCl, 2.5 KCl, 1.2 NaH₂PO₄, 30 NaHCO₃, 20 HEPES, 25 glucose, 5 sodium ascorbate, 3 sodium pyruvate, 5 N-acetylcysteine, 2 CaCl₂·4H₂O, and 2 MgSO₄·7H₂O (pH 7.30-7.32, osmolarity 299-301 mOsm). For recording, slices were transferred into a perfusion chamber where they were continuously perfused with oxygenated artificial cerebrospinal fluid (ACSF; 30-32°C) consisting of (in mM): 113 NaCl, 2.5 KCl, 1.2 MgSO_4_·7H_2_O, 2.5 CaCl_2_ ·2H_2_O, 1 NaH_2_PO_4_, 26 NaHCO_3_, 20 glucose, 3 sodium pyruvate, 1 sodium ascorbate (pH 7.30-7.32, osmolarity 299-301 mOsm), at a flow rate of 2-3ml/min.

All recordings were acquired using a Multiclamp 700B amplifier and digitized at a sampling rate of 10kHz using a Digidata 1440A A/D converter controlled by Clampex version 10.6 software (Axon Instruments, Union City, CA). Neurons were visualized using a Nikon microscope (Eclipse FN1, Nikon Instruments, Inc., Melville, NY) equipped with differential interference contrast microscopy. TdTomato-containing interneurons were identified using a series 120Q X-cite lamp (Excelitas Technologies, Waltham, MA) with an RFP filter. Recordings were obtained from layer V interneurons in the PL using borosilicate glass pipettes (3.5-6 MΩ). Liquid junction potentials were not accounted for in any experiment.

For whole cell current-clamp recordings (**Fig. 1-3**), pipettes were with filled with an intracellular solution containing (in mM) 125 K+-gluconate, 4 NaCl, 10 HEPES, 4 Mg-ATP, 0.3 Na-GTP, and 10 Na-phosphocreatine (pH 7.30-7.32, osmolarity 280-290 mOsm). Following break in, resting membrane potential was recorded for one minute. The cell was then transferred to current clamp and held at -70 mV where 600ms current injections were applied in 20pA increments from -100pA to 480pA to measure intrinsic and firing properties. For whole cell voltage-clamp recordings (**Fig. 4-6**), pipettes were filled with an intracellular solution containing (in mM) 120 CsOH, 120 D-gluconic acid, 2.8 NaCl, 20 HEPES, 5 TEA-Cl, 2.5 Mg-ATP, and 0.25 Na-GTP (pH 7.30-7.32, osmolarity 280-290 mOsm). Following break in, the cell was allowed to rest at -60mV for one minute prior to recording spontaneous excitatory postsynaptic currents (sEPSCs) at -60 mV and then was allowed to adjust to +10 mV for two minutes before recording spontaneous inhibitory postsynaptic currents (sIPSCs).

**Figure 1:**
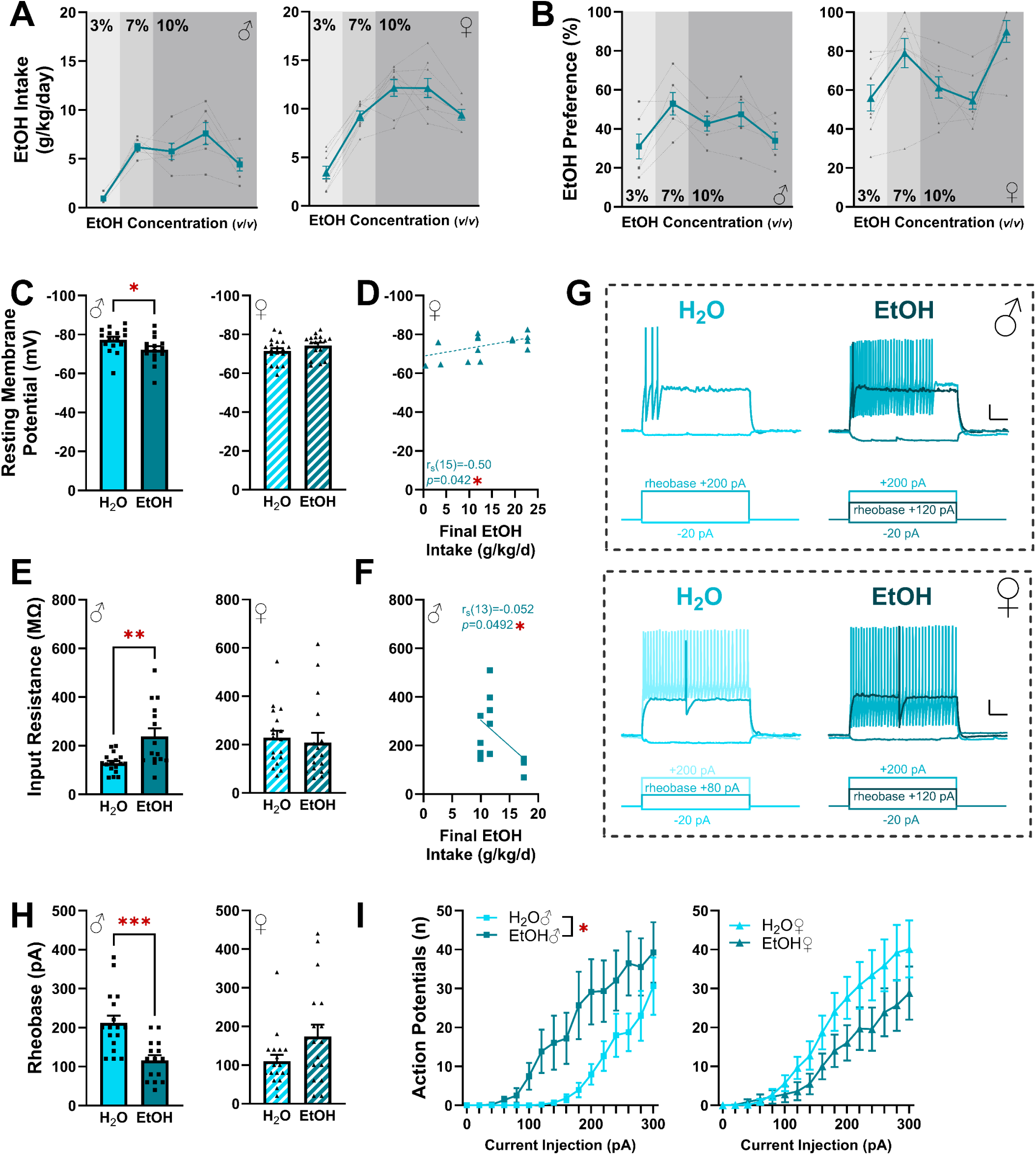
Effects of EtOH on intrinsic properties of prelimbic parvalbumin (PV) interneurons. Male data is shown on the left and female data on the right for all panels unless otherwise noted. **A**: EtOH intake in PV:Ai14 mice. **B**: EtOH preference in male PV:Ai14 mice. **C**: EtOH withdrawal depolarized PV interneurons in males. **D**: In females, EtOH intake was inversely correlated with resting membrane potential. **E:** EtOH withdrawal increased input resistance in males. **F**: Increased EtOH intake was correlated with decreased input resistance in males. **G**: Representative traces from current clamp recordings shown in H-I; scale bars represent 10mV, 100ms. **H:** EtOH withdrawal reduced rheobase in males. **I**: EtOH withdrawal increased action potential firing in males. \**p*<0.05, \*\**p*<0.01, \*\*\**p*<0.001. Data are shown as individual points where applicable, with bars representing the mean ± SEM when relevant. Specific sample sizes and statistical analyses can be found in Table 1.

**Table 1:** Statistical Analyses

| Inter-neuron Type | Figure Panel |  | Sex | Figure Description | Sample Size | Normality Test Results | Statistical Analysis Method | Test Statistic & *p*-value |
| --- | --- | --- | --- | --- | --- | --- | --- | --- |
| PV | 1 | C | Males | Resting Membrane Potential | H<sub>2</sub>O: 16 cells/5 mice<br>EtOH: 15 cells/3 mice | H<sub>2</sub>O: W=0.9110, *p*=0.1207; EtOH: W=0.9620, *p*=0.7273 | unpaired t test | t(29)=2.137, *p*=0.0411 |
| PV | 1 | C | Females | Resting Membrane Potential | H<sub>2</sub>O: 19 cells/6 mice<br>EtOH: 17 cells/7 mice | H<sub>2</sub>O: W=0.9609, *p*=0.5903; EtOH: W=0.8734, *p*=0.0249 | Mann Whitney U test | U=107, *p*=0.0872 |
| PV | 1 | D | Females | Resting Membrane Potential Correlation | 17 cells/7 mice | EtOH intake: K<sup>2</sup>=2.748, *p*=0.2531 | Spearman correlation | r<sub>s</sub>(15)=-0.50004, *p*=0.0426 |
| PV | 1 | F | Males | Input Resistance | H<sub>2</sub>O: 17 cells/5 mice<br>EtOH: 15 cells/3 mice | H<sub>2</sub>O: W=0.9476, *p*=0.4203; EtOH: W=0.8947, *p*=0.0791 | Welch's t test | t(16.33)=3.181, *p*=0.0057 |
| PV | 1 | F | Females | Input Resistance | H<sub>2</sub>O: 18 cells/6 mice<br>EtOH: 17cells/6mice | H<sub>2</sub>O: W=0.9303, *p*=0.1963; EtOH: W=0.7961, *p*=0.0018 | Mann Whitney U test | U=116, *p*=0.2315 |
| PV | 1 | G | Males | Input Resistance Correlation | 15 cells/3 mice | EtOH intake: K<sup>2</sup>=6.6165, *p*=0.0458 | Spearman correlation | r<sub>s</sub>(13)=-0.05204, *p*=0.0492 |
| PV | 1 | H | Males | Rheobase | H<sub>2</sub>O: 17 cells/5 mice<br>EtOH: 15 cells/3 mice | H<sub>2</sub>O: K<sup>2</sup>=2.755, *p*=0.2522; EtOH: K<sup>2</sup>=1.256, *p*=9.5338 | unpaired t test | t(30)=4.036, *p*=0.0003 |
| PV | 1 | H | Females | Rheobase | H<sub>2</sub>O: 18 cells/6 mice<br>EtOH: 18 cells/7 mice | H<sub>2</sub>O: K<sup>2</sup>=21.13, *p*<0.0001; EtOH: K<sup>2</sup>=2.900, *p*=0.2346 | Mann Whitney U test | U=111.5, *p*=0.1107 |
| PV | 1 | I | Males | Frequency-Current Plot | H<sub>2</sub>O: 17cells/5mice<br>EtOH: 15 cells/3mice | H<sub>2</sub>O: K<sup>2</sup>=4.487, *p*=0.1061; EtOH: K<sup>2</sup>=6.512, *p*=0.0385 | 2 way ANOVA | Current Injection x EtOH: F(15,450)=1.944, *p*=0.0178 ; Current Injection: F(2.416, 72.49)=18.74, *p*<0.0001; EtOH: F(1,30)=6.925, *p*=0.0133 |
| PV | 1 | I | Females | Frequency-Current Plot | H<sub>2</sub>O: 19 cells/6 mice<br>EtOH: 18 cells/7 mice | H<sub>2</sub>O: K<sup>2</sup>=6.829, *p*=0.0329; EtOH: K<sup>2</sup>=3.848, *p*=0.1461 | 2 way ANOVA | Current Injection x EtOH: F(15,525)=1.626, *p*=0.0627 ; Current Injection: F(1.397, 48.56)=34.49, *p*<0.0001; EtOH: F(1,35)=16.40, *p*=0.1334 |
| SOM | 2 | C | Males | Resting Membrane Potential | H<sub>2</sub>O: 15 cells/5 mice<br>EtOH: 15 cells/4 mice | H<sub>2</sub>O: W=0.8889, *p*=0.0645; EtOH: W=0.9567, *p*=0.6531 | unpaired t test | t(28)=1.068, *p*=0.2946 |
| SOM | 2 | C | Females | Resting Membrane Potential | H<sub>2</sub>O: 16 cells/5 mice<br>EtOH: 19 cells/6mice | H<sub>2</sub>O: W=0.9181, *p*=0.1575; EtOH: W=0.9598, *p*=0.5675 | unpaired t test | t(33)=0.01594, *p*=0.9874 |
| SOM | 2 | D | Males | Input Resistance | H<sub>2</sub>O: 15 cells/5 mice<br>EtOH: 15 cells/4 mice | H<sub>2</sub>O: W=0.9755, *p*=0.9294; EtOH: W=0.8568, *p*=0.0217 | Mann Whitney U test | U=107, *p*=0.8381 |
| SOM | 2 | D | Females | Input Resistance | H<sub>2</sub>O: 16 cells/5 mice<br>EtOH: 19 cells/6 mice | H<sub>2</sub>O: W=0.8944, *p*=0.0655; EtOH: W=0.9482, *p*=0.3684 | unpaired t test | t(33)=0.3643, *p*=0.7180 |
| SOM | 2 | E | Females | Input Resistance Correlation | 19 cells/6 mice | EtOH Intake: K<sup>2</sup>=0.7527, *p*=0.6864 | Pearson correlation | r(17)=0.6028, *p*=0.0063 |
| SOM | 2 | G | Males | Rheobase | H<sub>2</sub>O: 15 cells/5 mice<br>EtOH: 15 cells/4 mice | H<sub>2</sub>O: K<sup>2</sup>=5.367, *p*=0.0682; EtOH: K<sup>2</sup>=2.269, *p*=0.3216 | Welch's t test | t(21.72)=0.08244, *p*=0.9351 |
| SOM | 2 | G | Females | Rheobase | H<sub>2</sub>O: 16 cells/5 mice<br>EtOH: 19 cells/6mice | H<sub>2</sub>O: K<sup>2</sup>=3.040, *p*=0.2198; EtOH: K<sup>2</sup>=3.939, *p*=0.1395 | Mann Whitney U test | U=116, *p*=0.2326 |
| SOM | 2 | H | Males | Frequency-Current Plot | H<sub>2</sub>O: 17 cells/8 mice<br>EtOH: 17 cells/8 mice | H<sub>2</sub>O: K<sup>2</sup>=5.530, *p*=0.0630; EtOH: K<sup>2</sup>=8.258, *p*=0.0161 | 2 way ANOVA | Current Injection x EtOH: F(10,420)=0.3365, *p*=0.9913; Current Injection: F(1.647,44.01)=37.83, *p*<0.0001; EtOH: F(1,28)=0.5066, *p*=0.02302 |
| SOM | 2 | H | Females | Frequency-Current Plot | H<sub>2</sub>O: 16 cells/5 mice<br>EtOH: 19 cells/6 mice | H<sub>2</sub>O: K<sup>2</sup>=4.006, *p*=0.1349; EtOH: K<sup>2</sup>=2.121, *p*=0.3463 | 2 way ANOVA | Current Injection x EtOH: F(16,528)=0.2914, *p*=0.9970; Current Injection: F(2.042, 67.39)=17.30, *p*<0.0001; EtOH: F(1,33)=0.6992, *p*=0.4091 |
| VIP | 3 | C | Males | Resting Membrane Potential | H<sub>2</sub>O: 17 cells/8 mice<br>EtOH: 17 cells/8 mice | H<sub>2</sub>O: W=0.8769, *p*=0.0284; EtOH: W=0.9610, *p*=0.6503 | Mann Whitney U test | U=111, *p*=0.2594 |
| VIP | 3 | C | Females | Resting Membrane Potential | H<sub>2</sub>O: 16 cells/6 mice<br>EtOH: 15 cells/5 mice | H<sub>2</sub>O: W=0.9235, *p*=0.1918; EtOH: W=0.8980, *p*=0.0888 | unpaired t test | t(29)=2.176, *p*=0.0378 |
| VIP | 3 | D | Females | Resting Membrane Potential Correlation | 15 cells/5 mice | EtOH intake: K<sup>2</sup>=4.167, *p*=0.1245 | Pearson correlation | r(13)=0.7285, *p*=0.0021 |
| VIP | 3 | F | Males | Input Resistance | H<sub>2</sub>O: 17 cells/8 mice<br>EtOH: 17 cells/8 mice | H<sub>2</sub>O: W=0.9637, *p*=0.7025; EtOH: W=0.9645, *p*=0.7172 | unpaired t test | t(32)=0.2281, *p*=0.8210 |
| VIP | 3 | F | Females | Input Resistance | H<sub>2</sub>O: 16 cells/6 mice<br>EtOH: 14 cells/5 mice | H<sub>2</sub>O: W=0.9570, *p*=0.6086; EtOH: W=0.9242, *p*=0.2527 | unpaired t test | t(28)=4.079, *p*=0.0003 |
| VIP | 3 | G | Females | Input Resistance Correlation | 14 cells/5 mice | EtOH intake: K<sup>2</sup>=3.234, *p*=0.1985 | Pearson correlation | r(12)=0.7685, *p*=0.0013, |
| VIP | 3 | H | Males | Rheobase | H<sub>2</sub>O: 16 cells/7 mice<br>EtOH: 16 cells/8 mice | H<sub>2</sub>O: K<sup>2</sup>=1.055, *p*=0.5900; EtOH: K<sup>2</sup>=3.702, *p*=0.1571 | unpaired t test | t(30)=1.199, *p*=0.2398 |
| VIP | 3 | H | Females | Rheobase | H<sub>2</sub>O: 16 cells/6 mice<br>EtOH: 15 cells/5 mice | H<sub>2</sub>O: K<sup>2</sup>=1.215, *p*=0.5447; EtOH: K<sup>2</sup>=2.784, *p*=0.2561 | unpaired t test | t(29)=0.7664, *p*=0.4496 |
| VIP | 3 | I | Males | Frequency-Current Plot | H<sub>2</sub>O: 17 cells/8 mice<br>EtOH: 17 cells/8 mice | H<sub>2</sub>O: K<sup>2</sup>=6.924, *p*=0.0314; EtOH: K<sup>2</sup>=2.626, *p*=0.2691 | 2 way ANOVA | Current Injection x EtOH: F(15,480)=0.6586, *p*=0.8254 ; Current Injection: F(2.410, 77.11)=5.742, *p*=0.0028; EtOH: F(1,32)=0.2519, *p*=0.6192 |
| VIP | 3 | I | Females | Frequency-Current Plot | H<sub>2</sub>O: 16 cells/6 mice<br>EtOH: 15 cells/5 mice | H<sub>2</sub>O: K<sup>2</sup>=2.373, *p*=0.3052; EtOH: K<sup>2</sup>=4.885, *p*=0.0870 | 2 way ANOVA | Current Injection x EtOH: F(15,435)=0.8727, *p*=0.5954; Current Injection: F(1.974, 57.25)=12.25, *p*<0.0001; EtOH: F(1,29)=0.05451, *p*=0.8170 |
| PV | 4 | A | Males | sEPSC Frequency | H<sub>2</sub>O: 14 cells/5 mice<br>EtOH: 13 cells/3 mice | H<sub>2</sub>O: W=0.9256, *p*=0.2648; EtOH: W=0.8888, *p*=0.0940 | unpaired t test | t(25)=1.234, *p*=0.2287 |
| PV | 4 | A | Females | sEPSC Frequency | H<sub>2</sub>O: 13 cells/6 mice<br>EtOH: 12 cells/6 mice | H<sub>2</sub>O: W=0.9382, *p*=0.4335; EtOH: W=0.8869, *p*=0.1074 | unpaired t test | t(23)=0.07227, *p*=0.9430 |
| PV | 4 | B | Males | sEPSC amplitude | H<sub>2</sub>O: 15cells/5mice<br>EtOH: 13 cells/3mice | H<sub>2</sub>O: W=0.8974, *p*=0.0868; EtOH: W=0.9840, *p*=0.9934 | Welch's t test | t(19.23)=0.7100, *p*=0.4870 |
| PV | 4 | B | Females | sEPSC amplitude | H<sub>2</sub>O: 14 cells/6 mice<br>EtOH: 12 cells/6 mice | H<sub>2</sub>O: W=0.9563, *p*=0.6624; EtOH: W=0.9353, *p*=0.4394 | unpaired t test | t(24)=1.174, *p*=0.2520 |
| PV | 4 | D | Males | sIPSC Frequency | H<sub>2</sub>O: 15 cells/5 mice<br>EtOH: 13 cells/3 mice | H<sub>2</sub>O: W=0.8971, *p*=0.0859; EtOH: W=0.9123, *p*=0.1969 | t test with Welch's | t(15.94)=2.751, *p*=0.0142 |
| **PV** | 4 | D | Females | sIPSC Frequency | H<sub>2</sub>O: 13 cells/6 mice<br>EtOH: 13 cells/7 mice | H<sub>2</sub>O: W=0.9563, *p*=0.6624; EtOH: W=0.6733, *p*=0.0003 | Mann Whitney U test | U=61, *p*=0.5114 |
| **PV** | 4 | E | Males | sIPSC amplitude | H<sub>2</sub>O: 15cells/5mice<br>EtOH: 13 cells/3mice | H<sub>2</sub>O: W=0.9682, *p*=0.8313; EtOH: W=0.9131, *p*=0.2019 | unpaired t test | t(26)=0.5717, *p*=0.5725 |
| **PV** | 4 | E | Females | sIPSC amplitude | H<sub>2</sub>O: 13 cells/6 mice<br>EtOH: 13 cells/7 mice | H<sub>2</sub>O: W=0.9147, *p*=0.2125; EtOH: W=0.9084, *p*=0.4414 | unpaired t test | t(24)=2.944, *p*=0.0041 |
| **PV** | 4 | G | Males | E/I Ratio | H<sub>2</sub>O: 14cells/5mice<br>EtOH: 13 cells/3mice | H<sub>2</sub>O: W=0.5911, *p*<0.0001; EtOH: W=0.7926, *p*=0.0056 | Mann Whitney U test | U=36, *p*=0.0066 |
| **PV** | 4 | G | Females | E/I Ratio | H<sub>2</sub>O: 13 cells/6 mice<br>EtOH: 12 cells/6 mice | H<sub>2</sub>O: W=0.9263, *p*=0.3048; EtOH: W=0.7160, *p*=0.0012 | Mann Whitney U test | U=66, *p*=0.5382 |
| **PV** | 4 | H | Males | E/I Ratio Correlation | 13 cells/3mice | EtOH intake: K<sup>2</sup>=8.825, *p*=0.0121 | Spearman correlation | *r*<sub>s</sub>(11)=-0.5795, *p*=0.0415 |
| **PV** | 4 | I | Males | Synaptic Drive | H<sub>2</sub>O: 14cells/5mice<br>EtOH: 14 cells/3mice | H<sub>2</sub>O: W=0.8046, *p*=0.0057; EtOH: W=0.7355, *p*=0.0013 | Mann Whitney U test | U=51, *p*=0.0543 |
| **PV** | 4 | I | Females | Synaptic Drive | H<sub>2</sub>O: 14 cells/6 mice<br>EtOH: 12 cells/6 mice | H<sub>2</sub>O: W=0.8985, *p*=0.1972; EtOH: W=0.6402, *p*=0.0002 | Mann Whitney U test | U=70, *p*=0.4940 |
| **SOM** | 5 | A | Males | sEPSC Frequency | H<sub>2</sub>O: 15 cells/5 mice<br>EtOH: 15 cells/4 mice | H<sub>2</sub>O: W=0.7749, *p*=0.0018; EtOH: W=0.8367, *p*=0.0113 | Mann Whitney U test | U=70, *p*=0.1736 |
| **SOM** | 5 | A | Females | sEPSC Frequency | H<sub>2</sub>O: 17cells/5 mice<br>EtOH: 18 cells/6mice | H<sub>2</sub>O: W=0.9407, *p*=0.3265; EtOH: W=0.8837, *p*=0.0301 | Mann Whitney U test | U=75, *p*=0.0093 |
| **SOM** | 5 | B | Males | sEPSC Amplitude | H<sub>2</sub>O: 16 cells/5 mice<br>EtOH: 14 cells/4 mice | H<sub>2</sub>O: W=0.9657, *p*=0.7652; EtOH: W=0.8797, *p*=0.0576 | unpaired t test | t(28)=0.4046, *p*=0.6888 |
| **SOM** | 5 | B | Females | sEPSC Amplitude | H<sub>2</sub>O: 17cells/5 mice<br>EtOH: 17 cells/6mice | H<sub>2</sub>O: W=0.9463, *p*=0.4005; EtOH: W=0.9540, *p*=0.5227 | unpaired t test | t(32)=0.1311, *p*=0.8965 |
| **SOM** | 5 | D | Males | sIPSC Frequency | H<sub>2</sub>O: 16 cells/5 mice<br>EtOH: 15 cells/4 mice | H<sub>2</sub>O: W=0.8788, *p*=0.0371; EtOH: W=0.8565, *p*=0.0215 | Mann Whitney U test | U=89, *p*=0.2316 |
| **SOM** | 5 | D | Females | sIPSC Frequency | H<sub>2</sub>O: 16 cells/5 mice<br>EtOH: 17 cells/6 mice | H<sub>2</sub>O: W=0.8677, *p*=0.0251; EtOH: W=0.8446, *p*=0.0089 | Mann Whitney U test | U=115.5, *p*=0.4710 |
| **SOM** | 5 | E | Males | sIPSC Amplitude | H<sub>2</sub>O: 15 cells/5 mice<br>EtOH: 15 cells/4 mice | H<sub>2</sub>O: W=0.8716, *p*=0.0356; EtOH: W=0.9643, *p*=0.7671 | Mann Whitney U test | U=81, *p*=0.2017 |
| **SOM** | 5 | E | Females | sIPSC Amplitude | H<sub>2</sub>O: 17 cells/5 mice<br>EtOH: 17 cells/6 mice | H<sub>2</sub>O: W=0.9575, *p*=0.5850; EtOH: W=0.9596, *p*=0.6242 | unpaired t test | t(32)=0.04447, *p*=0.9648 |
| **SOM** | 5 | G | Males | E/I Ratio | H<sub>2</sub>O: 15 cells/5 mice<br>EtOH: 14 cells/4 mice | H<sub>2</sub>O: W=0.6139, *p*<0.0001; EtOH: W=0.5848, *p*<0.0001 | Mann Whitney U test | U=99, *p*=0.8132 |
| **SOM** | 5 | G | Females | E/I Ratio | H<sub>2</sub>O: 17cells/5 mice<br>EtOH: 17 cells/6mice | H<sub>2</sub>O: W=0.8339, *p*=0.0062; EtOH: W=0/8241, *p*=0.0044 | Mann Whitney U test | U=79, *p*=0.0238 |
| **SOM** | 5 | H | Males | Synaptic Drive | H<sub>2</sub>O: 15 cells/5 mice<br>EtOH: 14 cells/4 mice | H<sub>2</sub>O: W=0.5821, *p*<0.0001; EtOH: W=0.6628, *p*=0.0002 | Mann Whitney U test | U=104, *p*=0.9829 |
| **SOM** | 5 | H | Females | Synaptic Drive | H<sub>2</sub>O: 17cells/5 mice<br>EtOH: 17 cells/6mice | H<sub>2</sub>O: W=0.8096, *p*=0.0028; EtOH: W=0.8181, *p*=0.0036 | Mann Whitney U test | U=77, *p*=0.0196 |
| **VIP** | 6 | A | Males | sEPSC Frequency | H<sub>2</sub>O: 13 cells/6 mice<br>EtOH: 16 cells/9 mice | H<sub>2</sub>O: W=0.7508, *p*=0.0019; EtOH: W=0.9078, *p*=0.1074 | Mann Whitney U test | U=50, *p*=0.0172 |
| **VIP** | 6 | A | Females | sEPSC Frequency | H<sub>2</sub>O: 20 cells/7 mice<br>EtOH: 14 cells/6 mice | H<sub>2</sub>O: W=0.7841, *p*=0.0005; EtOH: W=0.7565, *p*=0.0015 | Mann Whitney | U=137, *p*=0.9313 |
| **VIP** | 6 | B | Males | sEPSC Amplitude | H<sub>2</sub>O: 13 cells/6 mice<br>EtOH: 17 cells/9 mice | H<sub>2</sub>O: W=0.9289, *p*=0.3292; EtOH: W=0.9397, *p*=0.3145 | unpaired t test | t(28)=0.01300, *p*=0.9897 |
| **VIP** | 6 | B | Females | sEPSC Amplitude | H<sub>2</sub>O: 21cells/7 mice<br>EtOH: 15cells/7 mice | H<sub>2</sub>O: W=0.9578, *p*=0.4721; EtOH: W=0.9050, *p*=0.1134 | unpaired t test | t(34)=1.720, *p*=0.0946 |
| **VIP** | 6 | D | Males | sIPSC Frequency | H<sub>2</sub>O: 13 cells/6 mice<br>EtOH: 17 cells/9 mice | H<sub>2</sub>O: W=0.9248, *p*=0.2909; EtOH: W=0.8728, *p*=0.0243 | Mann Whitney U test | U=92, *p*=0.2814 |
| **VIP** | 6 | D | Females | sIPSC Frequency | H<sub>2</sub>O: 21cells/7 mice<br>EtOH: 14cells/7 mice | H<sub>2</sub>O: W=0.8413, *p*=0.0030; EtOH: W=0.8482, *p*=0.0210 | Mann Whitney U test | U=112, *p*=0.2456 |
| VIP | 6 | E | Males | sIPSC Amplitude | H<sub>2</sub>O: 13 cells/6 mice<br>EtOH: 17 cells/9 mice | H<sub>2</sub>O: W=0.9289, *p*=0.3292; EtOH: W=0.9397, *p*=0.3145 | unpaired t test | t(28)=0.013, *p*=0.9897 |
| VIP | 6 | E | Females | sIPSC Amplitude | H<sub>2</sub>O: 21cells/7 mice,<br>EtOH: 15cells/7 mice | H<sub>2</sub>O: W=0.9785, *p*=0.9024; EtOH: W=0.8498, *p*=0.0173 | Mann Whitney U test | U=134, *p*=0.4654 |
| VIP | 6 | G | Males | E/I Ratio | H<sub>2</sub>O: 13 cells/6 mice<br>EtOH: 17 cells/9 mice | H<sub>2</sub>O: W=0.8458, *p*=0.0252; EtOH: W=0.8210, *p*=0.0056 | Mann Whitney U test | U=45, *p*=0.0052 |
| VIP | 6 | G | Females | E/I Ratio | H<sub>2</sub>O: 20cells/7 mice<br>EtOH: 15cells/7 mice | H<sub>2</sub>O: W=0.7268, *p*<0.0001; EtOH: W=0.6825, *p*=0.0002 | Mann Whitney U test | U=132, *p*=0.5644 |
| VIP | 6 | H | Males | Synaptic Drive | H<sub>2</sub>O: 13 cells/6 mice<br>EtOH: 17 cells/9 mice | H<sub>2</sub>O: W=0.8496, *p*=0.0282; EtOH: W=0.8193, *p*=0.0038 | Mann Whitney U test | U=45, *p*=0.0052 |
| VIP | 6 | K | Females | Synaptic Drive | H<sub>2</sub>O: 20 cells/7 mice<br>EtOH: 14cells/6 mice | H<sub>2</sub>O: W=0.6436, *p*<0.0001; EtOH: W=0.6622, *p*=0.0002 | Mann Whitney U test | U=102, *p*=0.1922 |

Resting membrane potential and input resistance were calculated using ClampFit 10 (Molecular Devices, San Jose, CA) whereas firing properties were analyzed with StimFit^90^. Rheobase was defined as the minimum current required to elicit an action potential. Voltage clamp recordings were analyzed with MiniAnalysis (Synaptosoft, Fort Lee, NJ, USA). E/I ratios were calculated as (sEPSC frequency/sIPSC frequency). Synaptic drive was calculated as (sEPSC frequency x sEPSC amplitude) / (sIPSC frequency x sIPSC amplitude).

### Statistics

All statistical analyses were conducted using GraphPad Prism version 10.3.1 (GraphPad Software, Boston, MA) with the threshold for significance set to *p* < 0.05. Data from at least three mice were used for each group, with 1-5 cells recorded per mouse. No more than 33% of the cells in each data set originated from a single mouse. Grubbs’s outlier test was conducted on each dataset individually. For any identified outliers with multiple measurements, each measure was assessed independently, and only the affected cell was excluded from that particular dataset. Normality was assessed using the D’Agostino-Pearson test for rheobase and firing rate, and the Shapiro-Wilk test for all other datasets. Firing rate plots were analyzed with two-way mixed ANOVAs, where current injection served as the within-subject factor and EtOH as the between subject factor. In all other datasets comparing water drinking mice to EtOH withdrawn mice, groups were compared using an unpaired t-test or a Mann Whitney U test depending on results of the normality tests. Welch’s correction was used for unpaired t tests when F-tests indicated unequal variances. For correlation analyses, Pearson or Spearman tests were used based on whether the data followed a normal distribution or not. Data are presented as individual data points with bars as mean ± standard error of the mean when applicable.

## RESULTS

These experiments examined the effects of EtOH withdrawal on PL interneuron physiology. To identify specific interneuron subtypes, we used transgenic reporter mice expressing tdTomato in each targeted interneuron subtype. Mice were randomly assigned to either a water (control) or EtOH group. Those in the EtOH group underwent chronic drinking using the CA2BC model, followed by a 72h withdrawal period before sacrifice for electrophysiological recordings. Although PV, SOM, and VIP interneurons are distributed across cortical layers in the mPFC^53, 66, 67, 77, 91, 92^, the current studies focused on layer V of PL, as it contains projections to subcortical regions involved in EtOH-associated behaviors^47, 60, 67, 93–96^. Full statistical analyses are provided in **Table 1**.

### EtOH withdrawal and PV interneuron excitability

Male and female PV:Ai14 mice consistently consumed 5-10g/kg/d of EtOH (**Fig. 1A**) and developed an EtOH preference (**Fig. 1B**), consistent with previous studies^85^. To assess the effects of chronic EtOH on intrinsic properties and excitability of PV interneurons, we performed whole-cell patch clamp electrophysiology 72h after removal of the EtOH bottle. Our findings revealed that 72h EtOH withdrawal depolarized the resting membrane potential in males, but not females (**Fig. 1C**).

Interestingly, increased EtOH intake correlated with a more hyperpolarized resting membrane potential in females (**Fig. 1D**), but no such correlation was found in males (**Fig. S1A**). We also observed that EtOH withdrawal increased input resistance in males, but had no effect in females (**Fig. 1E**). Furthermore, EtOH intake was inversely correlated with input resistance in males (**Fig. 1F)**, while no significant correlation was observed in in females (**Fig. S1B**). EtOH withdrawal reduced rheobase and increased firing frequency in males, with no effects observed in females (**Fig. 1G-I**). Rheobase was not correlated with EtOH intake in either sex (**Fig. S1C-D**). These results indicate that EtOH withdrawal increases PV interneuron excitability in males, but not females, highlighting potential sex-specific differences in the modulation of feed-forward inhibition.

### EtOH withdrawal and SOM interneuron excitability

SOM:Ai14 mice also consumed EtOH consistently (**Fig. 2A**) and developed EtOH preference (**Fig. 2B**). Unlike PV interneurons, SOM interneurons showed no significant changes in resting membrane potential following EtOH withdrawal (**Fig. 2C**) and there was no correlation between resting membrane potential and EtOH intake in either sex (**Fig. S2A-B**). We found no effect of EtOH withdrawal on input resistance **(Fig. 2D)**, though higher input resistance correlated with increased EtOH intake in females (**Fig. 2E**) but not males (**Fig. S2C**). Similarly, EtOH withdrawal had no significant effect on rheobase or firing rate (**Fig. 2F-I**), and EtOH intake did not correlate with rheobase in either sex (**Fig. S2D-E**). Collectively, these results suggest that EtOH withdrawal does not significantly impact SOM interneuron excitability.

**Figure 2:**
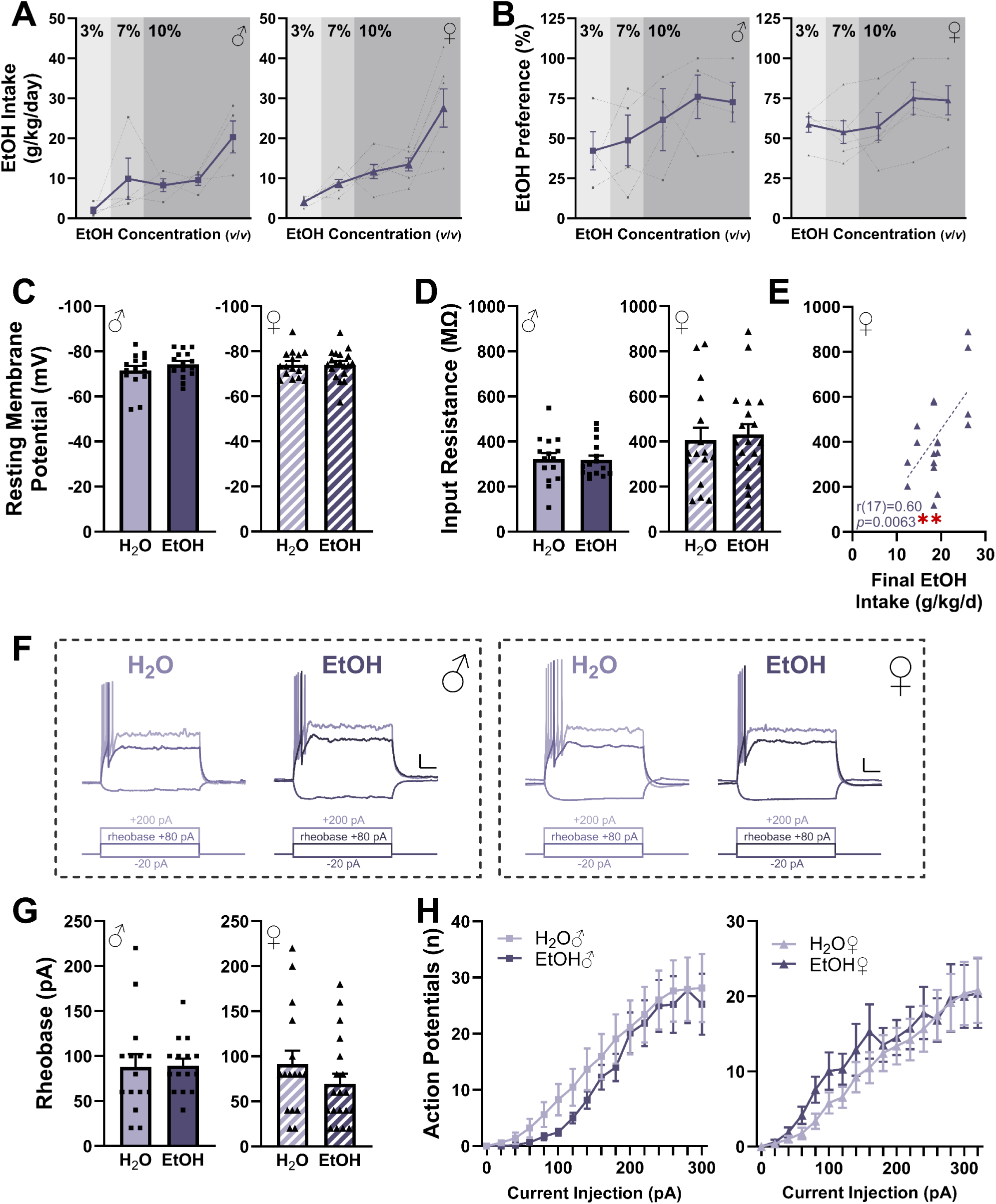
Effects of EtOH withdrawal on intrinsic properties of prelimbic somatostatin (SOM) interneurons. Male data is shown on the left and female data on the right for all panels unless otherwise noted. **A**: EtOH intake in SOM: Ai14 mice. **B**: EtOH preference in male SOM:Ai14 mice. **C**: Resting membrane potential in SOM interneurons was unaffected by EtOH withdrawal. **D**: Input resistance was unaffected by EtOH withdrawal. **E**: Increased EtOH intake was correlated with increased input resistance in females. **F:** Representative traces from current clamp recordings presented in G-H; scale bars represent 10mV, 100ms. **G:** Rheobase was unaffected by EtOH withdrawal. **H**: Action potential firing was unaffected by EtOH withdrawal. \*\**p*<0.01. Data are shown as individual points where applicable, with bars representing the mean ± SEM when relevant. Specific sample sizes and statistical analyses can be found in Table 1.

### EtOH withdrawal and VIP interneuron excitability

VIP:Ai14 mice consumed EtOH (**Fig. 3A**) and developed significant EtOH preference (**Fig. 3B**). In females, EtOH withdrawal hyperpolarized VIP interneurons (**Fig. 3C**), with effects primarily observed in low-drinking females (**Fig. 3D).** In contrast, EtOH withdrawal did not significantly affect resting membrane potential in males (**Fig. 3C**) and no correlation was found between resting membrane potential and EtOH intake (**Fig. S3A**). EtOH withdrawal reduced input resistance in females (**Fig. 3E),** which inversely correlated with EtOH consumption (**Fig. 3F**). No such changes were observed in males (**Fig. 3E, S3B)**. EtOH withdrawal did not significantly affect rheobase or firing rates (**Fig. 3G-I**) and EtOH intake did not correlate with rheobase in either sex (**Fig. S3C-D**). These findings suggest that EtOH withdrawal modulates VIP interneuron excitability in a sex-specific manner, with significant effects observed only in females.

**Figure 3:**
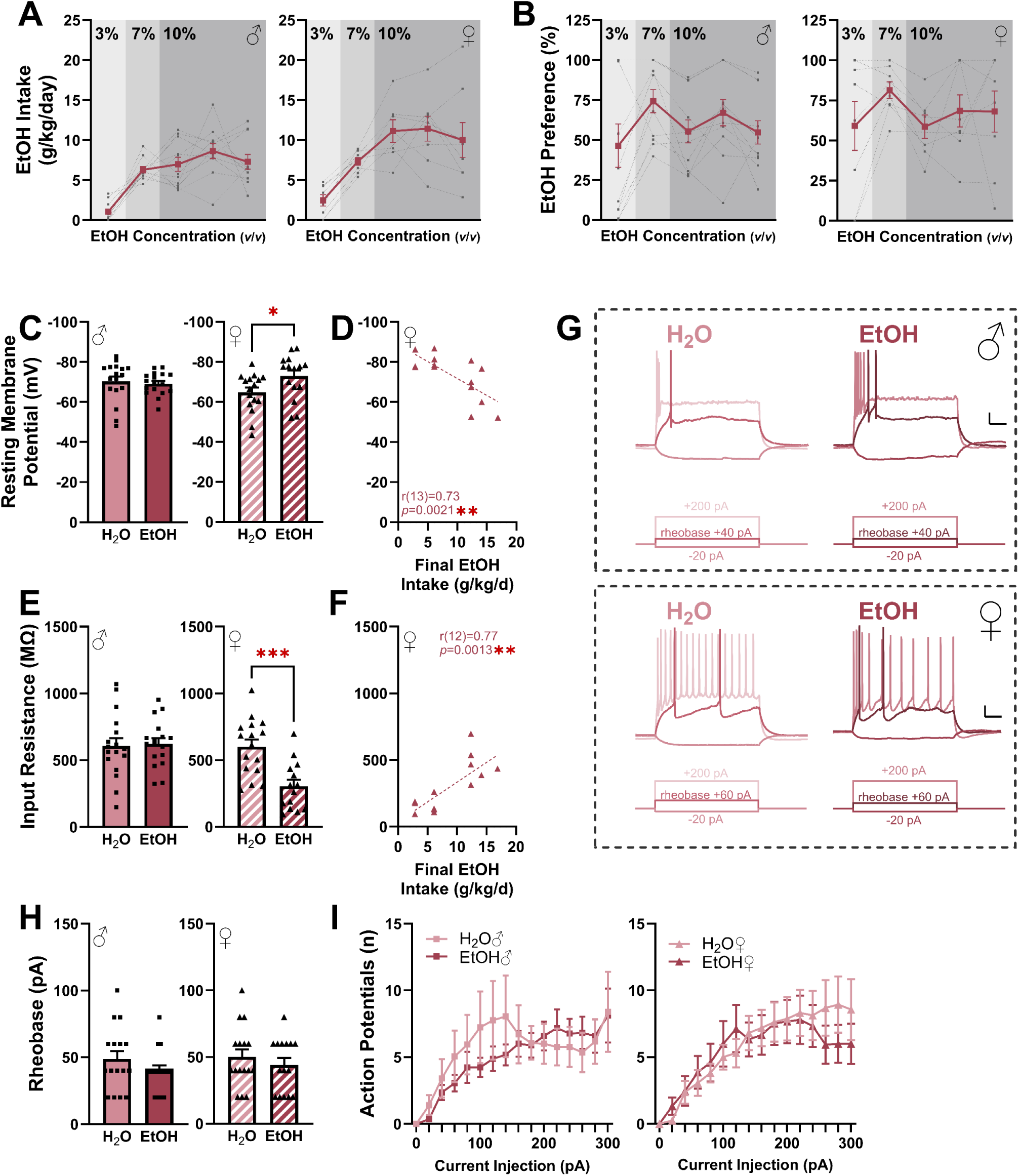
Effects of EtOH withdrawal on intrinsic properties of prelimbic vasoactive intestinal peptide (VIP) interneurons. Male data is shown on the left and female data on the right for all panels unless otherwise noted. **A**: EtOH intake in VIP:Ai14 mice. **B**: EtOH preference in male VIP:Ai14 mice. **C**: EtOH withdrawal hyperpolarized VIP interneurons in females. **D**: EtOH intake was correlated with resting membrane potential in females. **E:** EtOH withdrawal reduced input resistance in females. **F**: Increased EtOH intake was correlated with increased input resistance in females. **G**: Representative traces from current clamp recordings presented in H-I; scale bars represent 10mV, 100ms. **H:** Rheobase was unaffected by EtOH withdrawal. **I**: Action potential firing was unaffected by EtOH withdrawal. \**p*<0.05, ** *p*<0.01, ***p<0.001. Data are shown as individual points where applicable, with bars representing the mean ± SEM when relevant. Specific sample sizes and statistical analyses can be found in Table 1.

### EtOH withdrawal and PV interneuron synaptic transmission

To determine whether EtOH withdrawal affects synaptic input onto each interneuron subtype, we measured sEPSCs and sIPSCs while holding cells at -60 mV and +10 mV, respectively. In PV interneurons, no significant effects of EtOH withdrawal were observed on sESPC frequency or amplitude **(Fig. 4A-C).** However, EtOH withdrawal reduced sIPSC frequency in males (**Fig. 4D**), suggesting decreased GABAergic input onto PV interneurons in males. In contrast, sIPSC amplitude was increased in females following EtOH withdrawal (**Fig. 4E**; sIPSC traces in **Fig.4F**), while no significant changes were observed in males. EtOH intake did not correlate with frequency or amplitude of sEPSCs or sIPSCs in either sex (**Fig. S4A-H**).

**Figure 4:**
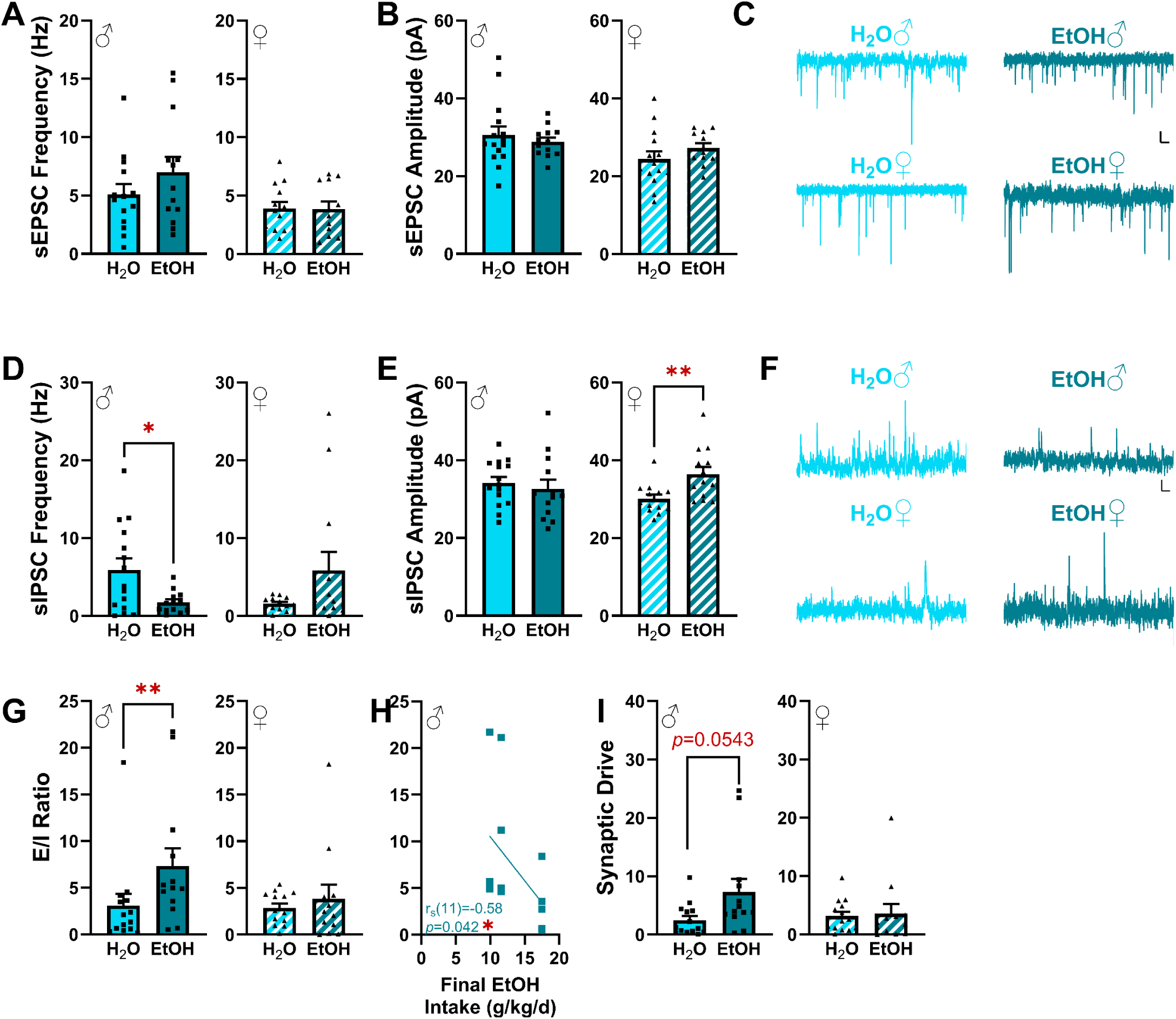
Effects of EtOH withdrawal on synaptic inputs to prelimbic parvalbumin (PV) interneurons. Male data is shown on the left and female data on the right for all panels unless otherwise noted. **A**: sEPSC frequency was unaffected by EtOH withdrawal. **B**: sEPSC amplitude was unaffected by EtOH withdrawal **C**: Representative sEPSC traces; scale bars represent 10mV, 100ms. **E**: EtOH withdrawal reduced sIPSC frequency in PV interneurons in males. **E:** EtOH withdrawal increased sIPSC amplitude in PV interneurons in females. **F:** Representative sIPSC traces; scale bars represent 10mV, 100ms. **G:** EtOH withdrawal increased E/I ratio in PV interneurons in males. **H:** Increased EtOH intake was correlated with reduced E/I ratios in males. **K:** There was a trend toward EtOH withdrawal increasing synaptic drive onto PV in males. \**p*<0.05, ** *p*<0.01. Data are shown as individual points where applicable, with bars representing the mean ± SEM when relevant. Specific sample sizes and statistical analyses can be found in Table 1.

EtOH withdrawal increased the E/I ratios in males (**Fig. 4G)**, and higher EtOH intake was associated with lower E/I ratios in males (**Fig. 4H**), suggesting that the effects seen in **Fig. 4G** were driven primarily by low-drinking male mice. EtOH intake did not significantly correlate with E/I ratios in females (**Fig. S4I**). Moreover, while there was a trend (*p*=0.0543) toward increased synaptic drive in males (**Fig.4I**), this trend did not reach statistical significance. Synaptic drive did not correlate with EtOH withdrawal (**Fig. S4J-K**). These results suggest that EtOH withdrawal increases the excitatory tone of PV interneurons in males.

### EtOH withdrawal and SOM interneuron synaptic transmission

In SOM neurons, EtOH withdrawal reduced sEPSC frequency in females (**Fig. 5A**), suggesting reduced glutamate release onto SOM neurons. However, no significant effects were observed on sEPSC amplitude (**Fig. 5B**; sEPSC traces in **Fig. 5C)** or sIPSC frequency and amplitude in either sex (**Fig. 5D-E**, sIPSC traces in **Fig. 5F**). Additionally, EtOH withdrawal reduced both E/I ratios (**Fig. 5G**) and synaptic drive (**Fig. 5H**) in females. Furthermore, EtOH intake did not correlate with synaptic transmission in either sex (**Fig. S5A-L**). These results suggest that EtOH withdrawal reduces the excitatory tone of SOM interneurons in females.

**Figure 5:**
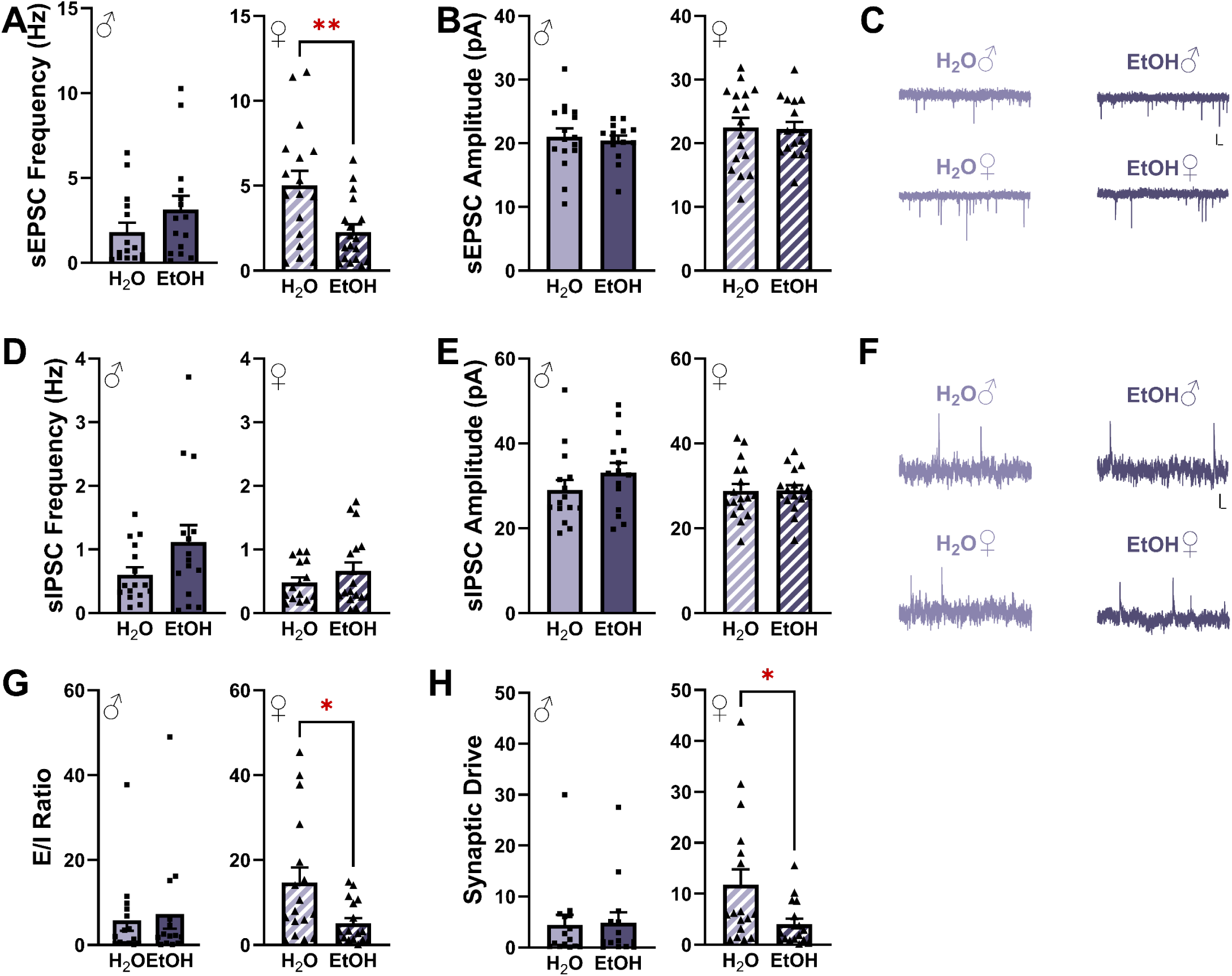
Effects of EtOH withdrawal on synaptic inputs to prelimbic somatostatin (SOM) interneurons. Male data is shown on the left and female data on the right for all panels unless otherwise noted. **A**: EtOH withdrawal increased sEPSC frequency in males. **B**: sEPSC amplitude was unaffected by EtOH withdrawal. **C**: Representative sEPSC traces; scale bars represent 10mV, 100ms. **D:** sIPSC frequency was unaffected by EtOH withdrawal. **E**: sIPSC amplitude was unaffected by EtOH withdrawal. **F**: Representative sIPSC traces; scale bars represent 10mV, 100ms. G: EtOH withdrawal increased E/I ratios in males. **H:** EtOH withdrawal increased synaptic drive onto SOM interneurons in males. *p<0.05, **p<0.01. Data are shown as individual points where applicable, with bars representing the mean ± SEM when relevant. Specific sample sizes and statistical analyses can be found in Table 1.

### EtOH withdrawal and VIP interneuron synaptic transmission

Finally, in VIP interneurons, EtOH withdrawal increased sEPSC frequency in males (**Fig. 6A**), suggesting enhanced glutamate release onto VIP interneurons in males. No significant changes were observed in sEPSC amplitude (**Fig. 6B**, sEPSC traces in **Fig. 6C**) or sIPSC frequency or amplitude (**Fig. 6D-E**, traces in **Fig. 6F**). EtOH withdrawal increased both the E/I ratios (**Fig. 6G**) and synaptic drive (**Fig. 6H**) in males. None of these parameters were correlated with EtOH intake (**Fig. S6A-L**). These results suggest that EtOH withdrawal increases excitatory input onto VIP interneurons in males.

**Figure 6:**
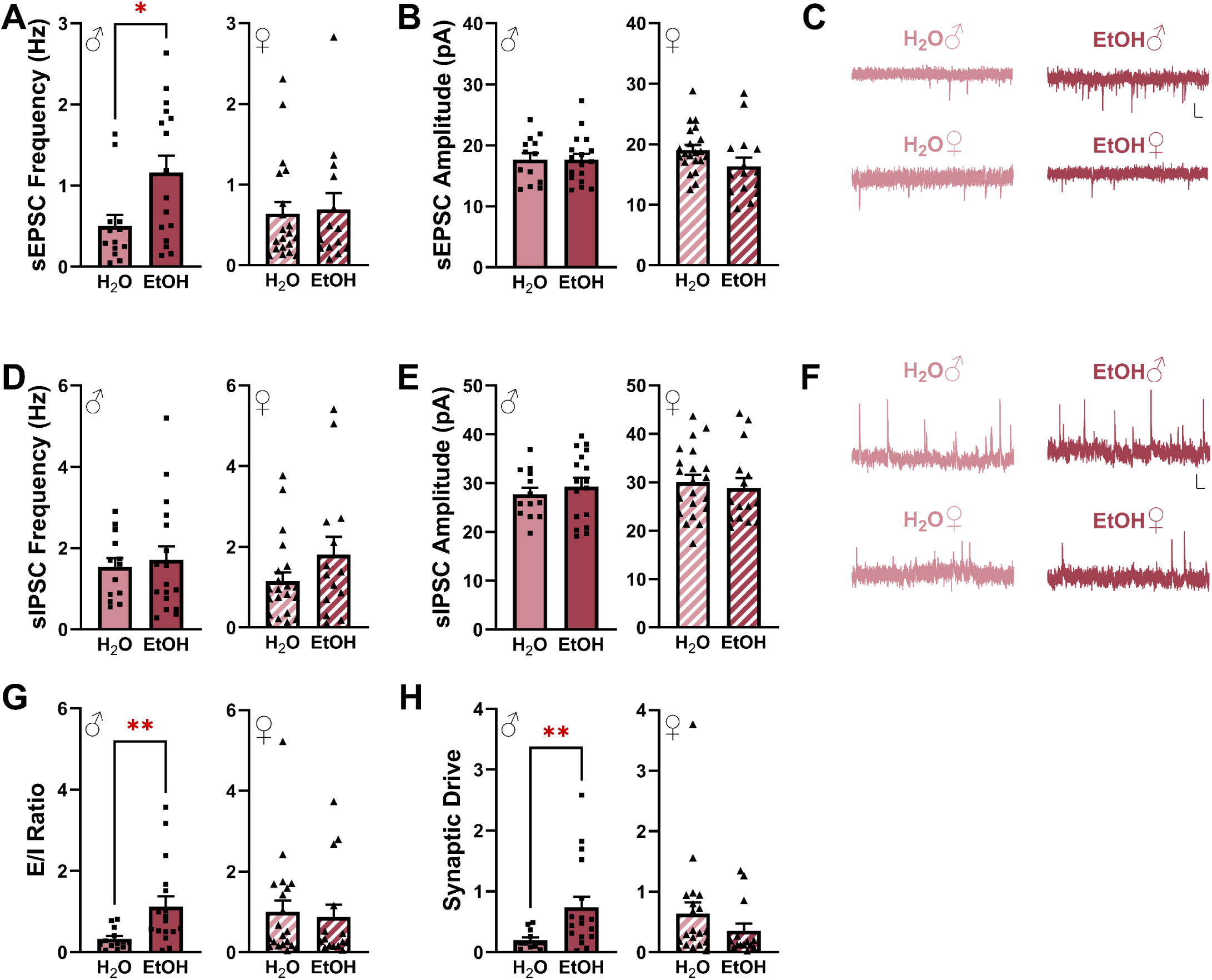
Effects of EtOH withdrawal on synaptic inputs to prelimbic vasoactive intestinal peptide (VIP) interneurons. Male data is shown on the left and female data on the right for all panels unless otherwise noted. **A**: EtOH withdrawal increased sEPSC frequency in males. **B**: sEPSC amplitude was unaffected by EtOH withdrawal. **C**: Representative sEPSC traces; scale bars represent 10mV, 100ms. **D**: sIPSC frequency was unaffected by EtOH withdrawal. **E:** sIPSC amplitude was unaffected by EtOH withdrawal. **F:** Representative sIPSC traces; scale bars represent 10mV, 100ms. **G:** EtOH withdrawal increased E/I ratios in males. **H:** EtOH withdrawal increased synaptic drive onto VIP interneurons in males. \**p*<0.05, \*\**p*<0.01. Data are shown as individual points where applicable, with bars representing the mean ± SEM when relevant. Specific sample sizes and statistical analyses can be found in Table 1.

A summary of all results is presented in **Table 2**.

**Table 2:** Summary of Findings

|  | PV |  |  |  | SOM+ |  |  |  | VIP+ |  |  |  |
| --- | --- | --- | --- | --- | --- | --- | --- | --- | --- | --- | --- | --- |
|  | Males |  | Females |  | Males |  | Females |  | Males |  | Females |  |
|  | group effects | correlation with EtOH intake | group effects | correlation with EtOH intake | group effects | correlation with EtOH intake | group effects | correlation with EtOH intake | group effects | correlation with EtOH intake | group effects | correlation with EtOH intake |
| RMP | EtOH more excitable | ns | ns | ↑ intake → ↓ excitability | ns | ns | ns | ns | ns | ns | EtOH decreased excitability | ↑ intake → ↑ excitability |
| Input Resistance | EtOH more excitable | ↑ intake → ↓ excitability | ns | ns | ns | ns | ns | ↑ intake → ↑ excitability | ns | ns | EtOH decreased excitability | ↑ intake → ↑ excitability |
| Rheobase | EtOH more excitable | ns | ns | ns | ns | ns | ns | ns | ns | ns | ns | ns |
| Firing Plot | EtOH more excitable | NA | ns | NA | ns | NA | ns | NA | ns | NA | ns | NA |
| sEPSC Frequency | ns | ns | ns | ns | ns | ns | EtOH decreased sEPSC frequency | ns | EtOH increased sEPSC frequency | ns | ns | ns |
| sEPSC Amplitude | ns | ns | ns | ↑ intake → ↑ amplitude | ns | ns | ns | ns | ns | ns | ns | ns |
| sIPSC Frequency | EtOH decreased sIPSC frequency | ns | ns | ↑ intake → ↑ frequency | ns | ns | ns | ns | ns | ns | ns | ns |
| sIPSC Amplitude | ns | ns | EtOH increased sIPSC amplitude | ns | ns | ns | ns | ns | ns | ns | ns | ns |
| E/I Frequency | EtOH increased E/I frequency | ns | ns | ns | ns | ns | EtOH decreased E/I frequency | ns | EtOH increased E/I frequency | ns | ns | ns |
| Synaptic Drive | trend toward EtOH increasing | ns | ns | ns | ns | ns | EtOH decreased synaptic drive | ns | EtOH increased synaptic drive | ns | ns | ns |

## DISCUSSION

Here we identified subtype- and sex-specific effects of EtOH withdrawal on interneuron excitability and synaptic physiology in layer V of PL. Specifically, EtOH withdrawal led to an increase excitability and a decrease in GABAergic inputs to PV interneurons, reduced glutamatergic inputs to SOM interneurons in females, and reduced excitability in VIP interneurons in females while increasing glutamatergic input in males. Furthermore, an increased EtOH intake correlated with heightened excitability in VIP and SOM neurons, whereas reduced EtOH intake was linked to decreased excitability in PV interneurons. Together these findings provide new insights into the impact of EtOH withdrawal on prelimbic inhibitory networks, underscoring the complexity of subtype- and sex-dependent synaptic adaptations.

Approximately 40% of all interneurons are PV interneurons^66^. Not only do PV interneurons contact nearly every pyramidal neuron nearby^97^, but the firing rate of PV interneurons increases as excitatory input stimuli increase^98^ suggesting PV interneurons mediate feedforward inhibition. Our results demonstrate EtOH withdrawal increases excitability of PV interneurons in male mice likely via increases in membrane resistance, although the precise ion channels underlying this effect remains to be determined. These data align with findings from a study using voluntary EtOH drinking models^51^; however, experimenter-administered models of EtOH exposure with shorter exposure periods (6-14d) have shown either reduced or no impact on excitability of PV interneurons^43, 52^. The longer exposures in our study and others (5+ weeks) suggest exposure duration and the nature of voluntary versus controlled administration paradigms may contribute to variability in PV interneuron response to EtOH withdrawal^51^. We also observed no change in glutamatergic synaptic transmission in PV interneurons following EtOH withdrawal, though previous studies have reported increased glutamate release^51^ or enhanced postsynaptic glutamatergic responses^51, 52^. Differences in withdrawal duration likely explain these discrepancies, as longer withdrawals are associated with amplified postsynaptic responses^52^. Notably, our study is among the first to examine EtOH’s withdrawal’s impact on GABAergic signaling onto PV interneurons. We found that EtOH withdrawal decreased GABA release in males but increased postsynaptic GABAergic responses in females.

While the reduced GABA release onto PV interneurons may ultimately contribute to the increased PV activity in males, the origin of the GABA reduction remains unclear. PV, SOM, and VIP interneurons each synapse onto PV interneurons^77^, yet EtOH withdrawal did not reduce excitability in these interneuron subtypes, suggesting that other interneuron populations may contribute to these effects.

SOM interneurons comprise approximately 30% of cortical interneurons and are involved primarily in feedback inhibition^66^. In the studies presented here, 72h EtOH withdrawal did not alter SOM excitability in either sex, although it reduced glutamate release onto SOM interneurons in females.

Reports on EtOH withdrawal’s effect on SOM interneuron synaptic plasticity are varied, likely due to differences in cortical layers examined, types of EtOH exposure, and withdrawal durations. For example, the drinking in the dark (DID) model reduced layer II/III SOM excitability in both sexes^33^ whereas intermittent access to 2 bottle choice (IA2BC) did not alter layer V SOM excitability in either sex^51^. Furthermore, EtOH reduced glutamate release in both sexes at 24h withdrawal in both the DID and IA2BC studies, suggesting the effects in males may resolve more quickly than in females^33, 51^.

While PV and SOM interneurons synapse onto and inhibit pyramidal neurons directly, VIP interneurons mainly synapse onto PV and SOM interneurons, thus disinhibiting pyramidal neurons^66^. To date, only one study has examined the effects of EtOH on VIP interneurons, finding that withdrawal increased excitability of layer II-III VIP interneurons in both sexes without altering excitatory synaptic transmission in either sex^99^. Indeed, most VIP interneurons are found in these superficial layers; however, a smaller proportion (approximately 10%) of VIP interneurons are located in deeper layers, which have not yet been explored in the context of EtOH exposure^53, 66, 67, 77, 91^. To our knowledge, this study is the first to investigate the effects of EtOH withdrawal on layer V VIP interneurons. Our results showed that EtOH withdrawal decreased VIP excitability in females, while increasing glutamate release onto these interneurons in males. Differences in drinking paradigms e.g. IA2BC vs CA2BC) or withdrawal length (24h vs 72h) may account for some of these findings. We hypothesize that layer-specific characteristics influence VIP interneurons’ response to EtOH withdrawal, as VIP neurons in upper and deeper layers exhibit distinct morphological, physiological and transcriptional profiles^91, 100, 101^, which could underlie these differential responses to EtOH.

In addition to examining group effects of EtOH withdrawal on interneuron physiology, we investigated whether final EtOH intake was correlated with excitability and synaptic transmission. Final EtOH intake was associated with reduced input resistance and increased GABA release onto PV interneurons in males, contrasting with group-level effects that indicated EtOH withdrawal increased excitability and decreased GABA release. Furthermore, the relationship between EtOH intake and excitability reversed in VIP interneurons in females: increased EtOH intake was associated with depolarized resting membrane potential and increased input resistance, whereas group effects indicated both measures decreased compared to water controls. These findings suggest that changes in excitability and synaptic transmission may be driven primarily by low drinking mice; however, increasing sample size of each group will be needed to properly determine whether the differences between low and high drinking mice are significant.

EtOH withdrawal is also associated with heightened pain states, negative affect, and deficits in executive function (reviewed in ^4, 102–106^). In the studies presented here, the CA2BC model of EtOH drinking was used partially because it reliably induces hyperalgesia and negative affect^85^. Thus, we hypothesized that similar mechanisms mediate EtOH withdrawal and neuropathic pain. Indeed, the spared nerve injury (SNI) model of neuropathic pain induces a variety of interneuron neuroadaptations. Specifically, SNI increases PV interneuron excitability in layer V in males^107^, decreases SOM interneuron excitability in layers II-III in females^107^, does not alter SOM interneuron excitability in layer V in either sex^107^, and reduces activity (in vivo) of VIP interneurons in both males and females^84^; paralleling a majority of the findings of studies of layer V interneurons presented here. One reason for the observed differences in VIP responses may be that the in vivo studies encompass both superficial and deep layers, whereas the studies presented here focused solely on layer V. Furthermore, inhibition of PV interneurons and activation of VIP interneurons reduce SNI-induced hyperalgesia^84, 108^, suggesting these interneurons are directly involved in mediating pain responses. Future studies will investigate the role of these interneurons in mediating responses to EtOH withdrawal associated hyperalgesia.

EtOH withdrawal is also associated with negative affect and deficits in executive function. Indeed, PL PV and SOM activity is associated with increased negative affect on variety of tests including the sucrose preference test, elevated plus maze, novelty suppressed feeding test, and forced swim test ^75, 78, 109, 110^. In contrast, VIP interneuron activity rises when the rodent is in anxiogenic environments (such as the open arm of the elevated plus maze)^83^. Furthermore, activation of PV and SOM interneurons increases impulsivity on a go/no-go task whereas activation of VIP interneurons reduces impulsivity^72, 111^. Combined with the results of the current study, these studies suggest shared neurobiological mechanisms across three different domains often associated with EtOH withdrawal: hyperalgesia, negative affect, and cognition. Additional studies are needed to clarify the opposing roles of VIP versus SOM and PV interneurons in mediating both negative affect and executive function and whether EtOH withdrawal affects them.

Finally, interneurons are often studied in the context of their influence on pyramidal neuron activity, whether through direct inhibition (PV, SOM) or disinhibition (VIP) ^81^. Notably, PV neurons exert the highest level of inhibitory control onto pyramidal neurons, followed by SOM, then VIP neurons^81^. One limitation of this study is we focused solely on interneurons, without investigating pyramidal neurons. However, there are a number of studies examining the effects of EtOH withdrawal on GABAergic transmission and excitability of mPFC pyramidal neurons, with results showing layer- and EtOH model-specific effects ^33, 37, 42, 44^. For example, oral gavage and intermittent EtOH drinking reduced GABA release onto layer V mPFC neurons^32, 45^, though this effect was absent following chronic intermittent vapor. Furthermore, while intermittent EtOH drinking did not alter layer V mPFC firing^47^, chronic intermittent vapor exposure reduced action potential firing^35^. Future studies will investigate the effects of CA2BC on layer V pyramidal neurons and determine whether the changes observed in interneurons contribute to the altered pyramidal neuron synaptic plasticity.

Together, our studies identified EtOH withdrawal-induced synaptic plasticity in three different types of interneurons. Furthermore, we highlight a range of synaptic adaptations that occur in one sex but not the other, ultimately highlighting the importance of including both males and females in animal models of AUD. Though studies examining gender differences in mPFC activity in adults with AUD are scarce, studies in adolescents and young adults report significant interactions of gender by EtOH use on frontocortical volume and thickness. Notably, adolescent girls with a history of EtOH use exhibit smaller grey matter volumes, increased cortical thickness, and reduced frontocortical activation during spatial memory tests. In contrast, adolescent boys with a history of EtOH use exhibit larger grey matter volume, reduced cortical thickness, and greater frontocortical activation during spatial memory tests ^112–115^. Furthermore, women with AUD perform significantly worse than men with AUD on cognitive tests^116–120^, which may result from AUD-induced mPFC dysfunction. These studies illustrate the need to further investigate gender differences in AUD. Whether FDA-approved medications for AUD differ in efficacy between genders remains unclear; however, women consistently experience more adverse effects from these drugs compared to men^121^. Acamprosate was the last drug approved for AUD, over twenty years ago (in 2004^122^) and AUD prevalence has since risen from 8.5 %^123^ to 10.9%^1^. With AUD affecting men and women differently, these findings emphasize the need to investigate gender-specific therapies that address both the neurobiological and clinical dimensions of the disorder.

## Author Contributions

SGQ: Conceptualization, Investigation, Formal Analysis, Visualization, Writing—Original Draft, Writing – Review & Editing

SDZ: Investigation, Writing – Review & Editing MGC: Investigation, Writing – Review & Editing

SP: Supervision, Conceptualization, Writing – Original Draft, Writing – Review & Editing, Funding Acquisition

## Acknowledgements

This work was supported by National Institute of Health grant AA026186 (SP). We thank Farhana Yasmin for her technical assistance in the beginning phase of these experiments.

## Notes

### Competing Interest Statement

The authors have declared no competing interest.

## References

1. NSDUH. 2023 National Survey on Drug Use and Health (NSDUH) 2023 NSDUH Section 2: Tobacco Product Use, Nicotine Vaping, and Alcohol Use Tables - .1 to 2.47; Section 5: Substance Use Disorder and Treatment Tables - 5.1 to 5.35].

2. Koob GF, Le Moal M. Drug addiction, dysregulation of reward, and allostasis. Neuropsychopharmacology : official publication of the American College of Neuropsychopharmacology. 2001;24(2):97–129. Epub 2000/12/20. doi: 10.1016/S0893-133X(00)00195-0. PubMed PMID: 11120394.

3. Koob GF, Le Moal M. Neurobiology of Addiction. Academic Press 2005.

4. Koob GF, Volkow ND. Neurobiology of addiction: a neurocircuitry analysis. The lancet Psychiatry. 2016;3(8):760–73. Epub 2016/08/01. doi: 10.1016/S2215-0366(16)00104-8. PubMed PMID: 27475769.

5. Kranzler HR, Soyka M. Diagnosis and Pharmacotherapy of Alcohol Use Disorder: A Review. Jama. 2018;320(8):815–24. Epub 2018/09/01. doi: 10.1001/jama.2018.11406. PubMed PMID: 30167705.

6. Herman MA, Quadir SG. Pharmacology of Alcohol Use. Elsevier; 2021.

7. Fairbanks J, Umbreit A, Kolla BP, Karpyak VM, Schneekloth TD, Loukianova LL, Sinha S. Evidence-Based Pharmacotherapies for Alcohol Use Disorder: Clinical Pearls. Mayo Clin Proc. 2020;95(9):1964–77. Epub 20200520. doi: 10.1016/j.mayocp.2020.01.030. PubMed PMID: 32446635.

8. Mason BJ, Heyser CJ. Alcohol Use Disorder: The Role of Medication in Recovery. Alcohol Res. 2021;41(1):07. Epub 20210603. doi: 10.35946/arcr.v41.1.07. PubMed PMID: 34113531; PMCID: PMC8184096.

9. Piazza NJ, Vrbka JL, Yeager RD. Telescoping of alcoholism in women alcoholics. Int J Addict. 1989;24(1):19–28. doi: 10.3109/10826088909047272. PubMed PMID: 2759762.

10. Randall CL, Roberts JS, Del Boca FK, Carroll KM, Connors GJ, Mattson ME. Telescoping of landmark events associated with drinking: a gender comparison. Journal of studies on alcohol. 1999;60(2):252–60. doi: 10.15288/jsa.1999.60.252. PubMed PMID: 10091964.

11. Greenfield SF, Back SE, Lawson K, Brady KT. Substance abuse in women. Psychiatr Clin North Am. 2010;33(2):339–55. doi: 10.1016/j.psc.2010.01.004. PubMed PMID: 20385341; PMCID: PMC3124962.

12. Mann K, Batra A, Gunthner A, Schroth G. Do women develop alcoholic brain damage more readily than men? Alcoholism, clinical and experimental research. 1992;16(6):1052–6. doi: 10.1111/j.1530-0277.1992.tb00698.x. PubMed PMID: 1471759.

13. Hommer D, Momenan R, Rawlings R, Ragan P, Williams W, Rio D, Eckardt M. Decreased corpus callosum size among alcoholic women. Arch Neurol. 1996;53(4):359–63. doi: 10.1001/archneur.1996.00550040099019. PubMed PMID: 8929159.

14. Hommer D, Momenan R, Kaiser E, Rawlings R. Evidence for a gender-related effect of alcoholism on brain volumes. The American journal of psychiatry. 2001;158(2):198–204. doi: 10.1176/appi.ajp.158.2.198. PubMed PMID: 11156801.

15. Mann K, Ackermann K, Croissant B, Mundle G, Nakovics H, Diehl A. Neuroimaging of gender differences in alcohol dependence: are women more vulnerable? Alcoholism, clinical and experimental research. 2005;29(5):896–901. doi: 10.1097/01.alc.0000164376.69978.6b. PubMed PMID: 15897736.

16. Towers EB, Williams IL, Qillawala EI, Rissman EF, Lynch WJ. Sex/Gender Differences in the Time-Course for the Development of Substance Use Disorder: A Focus on the Telescoping Effect. Pharmacological reviews. 2023;75(2):217–49. Epub 20221212. doi: 10.1124/pharmrev.121.000361. PubMed PMID: 36781217; PMCID: PMC9969523.

17. Holzhauer CG, Cucciare M, Epstein EE. Sex and Gender Effects in Recovery From Alcohol Use Disorder. Alcohol Res. 2020;40(3):03. Epub 20201119. doi: 10.35946/arcr.v40.3.03. PubMed PMID: 33224697; PMCID: PMC7668196.

18. NSDUH. 2013 National Survey on Drug Use and Health (NSDUH) 2013 Table 5.8A – Substance Dependence or Abuse in the Past Year among Persons Aged 18 or Older, by Demographic Characteristics: Numbers in Thousands, 2013 and 4]. Available from: https://www.samhsa.gov/data/sites/default/files/cbhsq-reports/NSDUHDetailedTabs2018R2/NSDUHDetTabsSect5pe2018.htm#tab5-4a.

19. Zhao Y, Skandali N, Bethlehem RAI, Voon V. Mesial Prefrontal Cortex and Alcohol Misuse: Dissociating Cross-sectional and Longitudinal Relationships in UK Biobank. Biological psychiatry. 2022;92(11):907–16. Epub 20220321. doi: 10.1016/j.biopsych.2022.03.008. PubMed PMID: 35589437.

20. Beck A, Wustenberg T, Genauck A, Wrase J, Schlagenhauf F, Smolka MN, Mann K, Heinz A. Effect of brain structure, brain function, and brain connectivity on relapse in alcohol-dependent patients. Archives of general psychiatry. 2012;69(8):842–52. doi: 10.1001/archgenpsychiatry.2011.2026. PubMed PMID: 22868938.

21. Grusser SM, Wrase J, Klein S, Hermann D, Smolka MN, Ruf M, Weber-Fahr W, Flor H, Mann K, Braus DF, Heinz A. Cue-induced activation of the striatum and medial prefrontal cortex is associated with subsequent relapse in abstinent alcoholics. Psychopharmacology. 2004;175(3):296–302. doi: 10.1007/s00213-004-1828-4. PubMed PMID: 15127179.

22. Vollstadt-Klein S, Loeber S, Kirsch M, Bach P, Richter A, Buhler M, von der Goltz C, Hermann D, Mann K, Kiefer F. Effects of cue-exposure treatment on neural cue reactivity in alcohol dependence: a randomized trial. Biological psychiatry. 2011;69(11):1060–6. Epub 20110203. doi: 10.1016/j.biopsych.2010.12.016. PubMed PMID: 21292243.

23. Blaine SK, Wemm S, Fogelman N, Lacadie C, Seo D, Scheinost D, Sinha R. Association of Prefrontal-Striatal Functional Pathology With Alcohol Abstinence Days at Treatment Initiation and Heavy Drinking After Treatment Initiation. The American journal of psychiatry. 2020;177(11):1048–59. Epub 20200828. doi: 10.1176/appi.ajp.2020.19070703. PubMed PMID: 32854534; PMCID: PMC7606814.

24. Seo D, Jia Z, Lacadie CM, Tsou KA, Bergquist K, Sinha R. Sex differences in neural responses to stress and alcohol context cues. Hum Brain Mapp. 2011;32(11):1998–2013. Epub 20101215. doi: 10.1002/hbm.21165. PubMed PMID: 21162046; PMCID: PMC3236497.

25. Pfefferbaum A, Sullivan EV, Rosenbloom MJ, Mathalon DH, Lim KO. A controlled study of cortical gray matter and ventricular changes in alcoholic men over a 5-year interval. Archives of general psychiatry. 1998;55(10):905–12. doi: 10.1001/archpsyc.55.10.905. PubMed PMID: 9783561.

26. Barbier E, Johnstone AL, Khomtchouk BB, Tapocik JD, Pitcairn C, Rehman F, Augier E, Borich A, Schank JR, Rienas CA, Van Booven DJ, Sun H, Natt D, Wahlestedt C, Heilig M. Dependence-induced increase of alcohol self-administration and compulsive drinking mediated by the histone methyltransferase PRDM2. Molecular psychiatry. 2017;22(12):1746–58. Epub 20160830. doi: 10.1038/mp.2016.131. PubMed PMID: 27573876; PMCID: PMC5677579.

27. Hashimoto JG, Gavin DP, Wiren KM, Crabbe JC, Guizzetti M. Prefrontal cortex expression of chromatin modifier genes in male WSP and WSR mice changes across ethanol dependence, withdrawal, and abstinence. Alcohol. 2017;60:83–94. Epub 20170314. doi: 10.1016/j.alcohol.2017.01.010. PubMed PMID: 28433423; PMCID: PMC5497775.

28. De Clerck M, Manguin M, Henkous N, d’Almeida MN, Beracochea D, Mons N. Chronic alcohol-induced long-lasting working memory deficits are associated with altered histone H3K9 dimethylation in the prefrontal cortex. Front Behav Neurosci. 2024;18:1354390. Epub 20240301. doi: 10.3389/fnbeh.2024.1354390. PubMed PMID: 38495426; PMCID: PMC10941761.

29. Crowley NA, Magee SN, Feng M, Jefferson SJ, Morris CJ, Dao NC, Brockway DF, Luscher B. Ketamine normalizes binge drinking-induced defects in glutamatergic synaptic transmission and ethanol drinking behavior in female but not male mice. Neuropharmacology. 2019;149:35–44. Epub 20190204. doi: 10.1016/j.neuropharm.2019.02.003. PubMed PMID: 30731135; PMCID: PMC6420859.

30. Kroener S, Mulholland PJ, New NN, Gass JT, Becker HC, Chandler LJ. Chronic alcohol exposure alters behavioral and synaptic plasticity of the rodent prefrontal cortex. PloS one. 2012;7(5):e37541. Epub 2012/06/06. doi: 10.1371/journal.pone.0037541. PubMed PMID: 22666364; PMCID: 3364267.

31. Siddiqi MT, Podder D, Pahng AR, Athanason AC, Nadav T, Cates-Gatto C, Kreifeldt M, Contet C, Roberts AJ, Edwards S, Roberto M, Varodayan FP. Prefrontal cortex glutamatergic adaptations in a mouse model of alcohol use disorder. Addict Neurosci. 2023;9. Epub 20231114. doi: 10.1016/j.addicn.2023.100137. PubMed PMID: 38152067; PMCID: PMC10752437.

32. Klenowski PM, Fogarty MJ, Shariff M, Belmer A, Bellingham MC, Bartlett SE. Increased Synaptic Excitation and Abnormal Dendritic Structure of Prefrontal Cortex Layer V Pyramidal Neurons following Prolonged Binge-Like Consumption of Ethanol. eNeuro. 2016;3(6). Epub 20161223. doi: 10.1523/ENEURO.0248-16.2016. PubMed PMID: 28032119; PMCID: PMC5179982.

33. Dao NC, Brockway DF, Suresh Nair M, Sicher AR, Crowley NA. Somatostatin neurons control an alcohol binge drinking prelimbic microcircuit in mice. Neuropsychopharmacology : official publication of the American College of Neuropsychopharmacology. 2021;46(11):1906–17. Epub 2021/06/12. doi: 10.1038/s41386-021-01050-1. PubMed PMID: 34112959; PMCID: PMC8429551.

34. Seif T, Chang SJ, Simms JA, Gibb SL, Dadgar J, Chen BT, Harvey BK, Ron D, Messing RO, Bonci A, Hopf FW. Cortical activation of accumbens hyperpolarization-active NMDARs mediates aversion-resistant alcohol intake. Nature neuroscience. 2013;16(8):1094–100. Epub 20130630. doi: 10.1038/nn.3445. PubMed PMID: 23817545; PMCID: PMC3939030.

35. Przybysz KR, Shillinglaw JE, Wheeler SR, Glover EJ. Chronic ethanol exposure produces long-lasting, subregion-specific physiological adaptations in RMTg-projecting mPFC neurons. Neuropharmacology. 2024;259:110098. Epub 20240806. doi: 10.1016/j.neuropharm.2024.110098. PubMed PMID: 39117106.

36. Domi A, Cadeddu D, Lucente E, Gobbo F, Edvardsson C, Petrella M, Jerlhag E, Ericson M, Soderpalm B, Adermark L. Pre- and postsynaptic signatures in the prelimbic cortex associated with “alcohol use disorder” in the rat. Neuropsychopharmacology : official publication of the American College of Neuropsychopharmacology. 2024. Epub 20240516. doi: 10.1038/s41386-024-01887-2. PubMed PMID: 38755284.

37. Pleil KE, Lowery-Gionta EG, Crowley NA, Li C, Marcinkiewcz CA, Rose JH, McCall NM, Maldonado-Devincci AM, Morrow AL, Jones SR, Kash TL. Effects of chronic ethanol exposure on neuronal function in the prefrontal cortex and extended amygdala. Neuropharmacology. 2015;99:735–49. Epub 20150716. doi: 10.1016/j.neuropharm.2015.06.017. PubMed PMID: 26188147; PMCID: PMC4781662.

38. Trantham-Davidson H, Burnett EJ, Gass JT, Lopez MF, Mulholland PJ, Centanni SW, Floresco SB, Chandler LJ. Chronic alcohol disrupts dopamine receptor activity and the cognitive function of the medial prefrontal cortex. The Journal of neuroscience : the official journal of the Society for Neuroscience. 2014;34(10):3706–18. doi: 10.1523/JNEUROSCI.0623-13.2014. PubMed PMID: 24599469; PMCID: PMC3942586.

39. Avchalumov Y, Oliver RJ, Trenet W, Heyer Osorno RE, Sibley BD, Purohit DC, Contet C, Roberto M, Woodward JJ, Mandyam CD. Chronic ethanol exposure differentially alters neuronal function in the medial prefrontal cortex and dentate gyrus. Neuropharmacology. 2021;185:108438. Epub 20201215. doi: 10.1016/j.neuropharm.2020.108438. PubMed PMID: 33333103; PMCID: PMC7927349.

40. Patel RR, Gandhi P, Spencer K, Salem NA, Erikson CM, Borgonetti V, Vlkolinsky R, Rodriguez L, Nadav T, Bajo M, Roberts AJ, Dayne Mayfield R, Roberto M. Functional and morphological adaptation of medial prefrontal corticotropin releasing factor receptor 1-expressing neurons in male mice following chronic ethanol exposure. Neurobiol Stress. 2024;31:100657. Epub 20240617. doi: 10.1016/j.ynstr.2024.100657. PubMed PMID: 38983690; PMCID: PMC11231756.

41. Hu W, Morris B, Carrasco A, Kroener S. Effects of acamprosate on attentional set-shifting and cellular function in the prefrontal cortex of chronic alcohol-exposed mice. Alcoholism, clinical and experimental research. 2015;39(6):953–61. Epub 20150423. doi: 10.1111/acer.12722. PubMed PMID: 25903298; PMCID: PMC10782929.

42. Patel RR, Wolfe SA, Borgonetti V, Gandhi PJ, Rodriguez L, Snyder AE, D’Ambrosio S, Bajo M, Domissy A, Head S, Contet C, Dayne Mayfield R, Roberts AJ, Roberto M. Ethanol withdrawal-induced adaptations in prefrontal corticotropin releasing factor receptor 1-expressing neurons regulate anxiety and conditioned rewarding effects of ethanol. Molecular psychiatry. 2022;27(8):3441–51. Epub 20220606. doi: 10.1038/s41380-022-01642-3. PubMed PMID: 35668157; PMCID: PMC9708587.

43. Hughes BA, Crofton EJ, O’Buckley TK, Herman MA, Morrow AL. Chronic ethanol exposure alters prelimbic prefrontal cortical Fast-Spiking and Martinotti interneuron function with differential sex specificity in rat brain. Neuropharmacology. 2020;162:107805. Epub 20191004. doi: 10.1016/j.neuropharm.2019.107805. PubMed PMID: 31589884; PMCID: PMC7027948.

44. Varodayan FP, Sidhu H, Kreifeldt M, Roberto M, Contet C. Morphological and functional evidence of increased excitatory signaling in the prelimbic cortex during ethanol withdrawal. Neuropharmacology. 2018;133:470–80. Epub 20180219. doi: 10.1016/j.neuropharm.2018.02.014. PubMed PMID: 29471053; PMCID: PMC5865397.

45. Hughes BA, Bohnsack JP, O’Buckley TK, Herman MA, Morrow AL. Chronic Ethanol Exposure and Withdrawal Impair Synaptic GABA(A) Receptor-Mediated Neurotransmission in Deep-Layer Prefrontal Cortex. Alcoholism, clinical and experimental research. 2019;43(5):822–32. Epub 20190418. doi: 10.1111/acer.14015. PubMed PMID: 30860602; PMCID: PMC6502689.

46. Cannady R, Nguyen T, Padula AE, Rinker JA, Lopez MF, Becker HC, Woodward JJ, Mulholland PJ. Interaction of chronic intermittent ethanol and repeated stress on structural and functional plasticity in the mouse medial prefrontal cortex. Neuropharmacology. 2021;182:108396. Epub 20201109. doi: 10.1016/j.neuropharm.2020.108396. PubMed PMID: 33181147; PMCID: PMC7942177.

47. Joffe ME, Winder DG, Conn PJ. Increased Synaptic Strength and mGlu(2/3) Receptor Plasticity on Mouse Prefrontal Cortex Intratelencephalic Pyramidal Cells Following Intermittent Access to Ethanol. Alcoholism, clinical and experimental research. 2021;45(3):518–29. Epub 20210209. doi: 10.1111/acer.14546. PubMed PMID: 33434325; PMCID: PMC7969423.

48. Dao NC, Suresh Nair M, Magee SN, Moyer JB, Sendao V, Brockway DF, Crowley NA. Forced Abstinence From Alcohol Induces Sex-Specific Depression-Like Behavioral and Neural Adaptations in Somatostatin Neurons in Cortical and Amygdalar Regions. Front Behav Neurosci. 2020;14:86. Epub 20200527. doi: 10.3389/fnbeh.2020.00086. PubMed PMID: 32536856; PMCID: PMC7266989.

49. Fabian CB, Jordan ND, Cole RH, Carley LG, Thompson SM, Seney ML, Joffe ME. Parvalbumin interneuron mGlu(5) receptors govern sex differences in prefrontal cortex physiology and binge drinking. Neuropsychopharmacology : official publication of the American College of Neuropsychopharmacology. 2024. Epub 2024/05/22. doi: 10.1038/s41386-024-01889-0. PubMed PMID: 38773314.

50. Thompson SM, Fabian CB, Ferranti AS, Joffe ME. Acute alcohol and chronic drinking bidirectionally regulate the excitability of prefrontal cortex vasoactive intestinal peptide interneurons. Neuropharmacology. 2023;238:109638. Epub 2023/07/24. doi: 10.1016/j.neuropharm.2023.109638. PubMed PMID: 37482180; PMCID: PMC10529784.

51. Joffe ME, Winder DG, Conn PJ. Contrasting sex-dependent adaptations to synaptic physiology and membrane properties of prefrontal cortex interneuron subtypes in a mouse model of binge drinking. Neuropharmacology. 2020;178:108126. Epub 2020/08/12. doi: 10.1016/j.neuropharm.2020.108126. PubMed PMID: 32781000; PMCID: PMC7544622.

52. Ferranti AS, Johnson KA, Winder DG, Conn PJ, Joffe ME. Prefrontal cortex parvalbumin interneurons exhibit decreased excitability and potentiated synaptic strength after ethanol reward learning. Alcohol. 2022;101:17–26. Epub 20220226. doi: 10.1016/j.alcohol.2022.02.003. PubMed PMID: 35227826; PMCID: PMC9117490.

53. Xu X, Roby KD, Callaway EM. Immunochemical characterization of inhibitory mouse cortical neurons: three chemically distinct classes of inhibitory cells. J Comp Neurol. 2010;518(3):389–404. doi: 10.1002/cne.22229. PubMed PMID: 19950390; PMCID: PMC2804902.

54. Ferguson BR, Gao WJ. PV Interneurons: Critical Regulators of E/I Balance for Prefrontal Cortex-Dependent Behavior and Psychiatric Disorders. Front Neural Circuits. 2018;12:37. Epub 20180516. doi: 10.3389/fncir.2018.00037. PubMed PMID: 29867371; PMCID: PMC5964203.

55. Crowley NA, Joffe ME. Developing breakthrough psychiatric treatments by modulating G protein-coupled receptors on prefrontal cortex somatostatin interneurons. Neuropsychopharmacology : official publication of the American College of Neuropsychopharmacology. 2022;47(1):389–90. Epub 2021/08/04. doi: 10.1038/s41386-021-01119-x. PubMed PMID: 34341496; PMCID: PMC8617253 financial relationships considered a conflict of interest.

56. Melon LC, Nasman JT, John AS, Mbonu K, Maguire JL. Interneuronal delta-GABA(A) receptors regulate binge drinking and are necessary for the behavioral effects of early withdrawal. Neuropsychopharmacology : official publication of the American College of Neuropsychopharmacology. 2019;44(2):425–34. Epub 20180728. doi: 10.1038/s41386-018-0164-z. PubMed PMID: 30089884; PMCID: PMC6300562.

57. Patton MS, Heckman M, Kim C, Mu C, Mathur BN. Compulsive alcohol consumption is regulated by dorsal striatum fast-spiking interneurons. Neuropsychopharmacology : official publication of the American College of Neuropsychopharmacology. 2021;46(2):351–9. Epub 20200714. doi: 10.1038/s41386-020-0766-0. PubMed PMID: 32663841; PMCID: PMC7852608.

58. Nahar L, Grant CA, Hewett C, Cortes D, Reker AN, Kang S, Choi DS, Nam HW. Regulation of Pv-specific interneurons in the medial prefrontal cortex and reward-seeking behaviors. Journal of neurochemistry. 2021;156(2):212–24. Epub 20200720. doi: 10.1111/jnc.15106. PubMed PMID: 32594517; PMCID: PMC7765736.

59. DiLeo A, Antonoudiou P, Ha S, Maguire JL. Sex Differences in the Alcohol-Mediated Modulation of BLA Network States. eNeuro. 2022;9(4). Epub 20220708. doi: 10.1523/ENEURO.0010-22.2022. PubMed PMID: 35788104; PMCID: PMC9275151.

60. Patton MS, Sheats SH, Siclair AN, Mathur BN. Alcohol potentiates multiple GABAergic inputs to dorsal striatum fast-spiking interneurons. Neuropharmacology. 2023;232:109527. Epub 20230401. doi: 10.1016/j.neuropharm.2023.109527. PubMed PMID: 37011784; PMCID: PMC10122715.

61. Patton MH, Roberts BM, Lovinger DM, Mathur BN. Ethanol Disinhibits Dorsolateral Striatal Medium Spiny Neurons Through Activation of A Presynaptic Delta Opioid Receptor. Neuropsychopharmacology : official publication of the American College of Neuropsychopharmacology. 2016;41(7):1831–40. Epub 20151211. doi: 10.1038/npp.2015.353. PubMed PMID: 26758662; PMCID: PMC4869052.

62. Andrade JP, Fernando PM, Madeira MD, Paula-Barbosa MM, Cadete-Leite A, Zimmer J. Effects of chronic alcohol consumption and withdrawal on the somatostatin-immunoreactive neurons of the rat hippocampal dentate hilus. Hippocampus. 1992;2(1):65–71. doi: 10.1002/hipo.450020109. PubMed PMID: 1364047.

63. Wadsworth HA, Anderson EQ, Williams BM, Ronstrom JW, Moen JK, Lee AM, McIntosh JM, Wu J, Yorgason JT, Steffensen SC. Role of alpha6-Nicotinic Receptors in Alcohol-Induced GABAergic Synaptic Transmission and Plasticity to Cholinergic Interneurons in the Nucleus Accumbens. Molecular neurobiology. 2023;60(6):3113–29. Epub 20230218. doi: 10.1007/s12035-023-03263-5. PubMed PMID: 36802012; PMCID: PMC10690621.

64. Diaz MR, Christian DT, Anderson NJ, McCool BA. Chronic ethanol and withdrawal differentially modulate lateral/basolateral amygdala paracapsular and local GABAergic synapses. The Journal of pharmacology and experimental therapeutics. 2011;337(1):162–70. Epub 20110105. doi: 10.1124/jpet.110.177121. PubMed PMID: 21209156; PMCID: PMC3063746.

65. Munoz B, Atwood BK. A novel inhibitory corticostriatal circuit that expresses mu opioid receptor-mediated synaptic plasticity. Neuropharmacology. 2023;240:109696. Epub 20230901. doi: 10.1016/j.neuropharm.2023.109696. PubMed PMID: 37659438; PMCID: PMC10591984.

66. Tremblay R, Lee S, Rudy B. GABAergic Interneurons in the Neocortex: From Cellular Properties to Circuits. Neuron. 2016;91(2):260–92. doi: 10.1016/j.neuron.2016.06.033. PubMed PMID: 27477017; PMCID: PMC4980915.

67. Anastasiades PG, Carter AG. Circuit organization of the rodent medial prefrontal cortex. Trends in neurosciences. 2021;44(7):550–63. Epub 20210507. doi: 10.1016/j.tins.2021.03.006. PubMed PMID: 33972100; PMCID: PMC8222144.

68. Atallah BV, Bruns W, Carandini M, Scanziani M. Parvalbumin-expressing interneurons linearly transform cortical responses to visual stimuli. Neuron. 2012;73(1):159–70. doi: 10.1016/j.neuron.2011.12.013. PubMed PMID: 22243754; PMCID: PMC3743079.

69. Sparta DR, Hovelso N, Mason AO, Kantak PA, Ung RL, Decot HK, Stuber GD. Activation of prefrontal cortical parvalbumin interneurons facilitates extinction of reward-seeking behavior. The Journal of neuroscience : the official journal of the Society for Neuroscience. 2014;34(10):3699–705. doi: 10.1523/JNEUROSCI.0235-13.2014. PubMed PMID: 24599468; PMCID: PMC3942585.

70. Brown JA, Ramikie TS, Schmidt MJ, Baldi R, Garbett K, Everheart MG, Warren LE, Gellert L, Horvath S, Patel S, Mirnics K. Inhibition of parvalbumin-expressing interneurons results in complex behavioral changes. Molecular psychiatry. 2015;20(12):1499–507. Epub 20150127. doi: 10.1038/mp.2014.192. PubMed PMID: 25623945; PMCID: PMC4516717.

71. Kim H, Ahrlund-Richter S, Wang X, Deisseroth K, Carlen M. Prefrontal Parvalbumin Neurons in Control of Attention. Cell. 2016;164(1-2):208–18. doi: 10.1016/j.cell.2015.11.038. PubMed PMID: 26771492; PMCID: PMC4715187.

72. Kamigaki T, Dan Y. Delay activity of specific prefrontal interneuron subtypes modulates memory-guided behavior. Nature neuroscience. 2017;20(6):854–63. Epub 20170424. doi: 10.1038/nn.4554. PubMed PMID: 28436982; PMCID: PMC5554301.

73. Shepard R, Coutellier L. Changes in the Prefrontal Glutamatergic and Parvalbumin Systems of Mice Exposed to Unpredictable Chronic Stress. Molecular neurobiology. 2018;55(3):2591–602. Epub 20170418. doi: 10.1007/s12035-017-0528-0. PubMed PMID: 28421533.

74. Xu H, Liu L, Tian Y, Wang J, Li J, Zheng J, Zhao H, He M, Xu TL, Duan S, Xu H. A Disinhibitory Microcircuit Mediates Conditioned Social Fear in the Prefrontal Cortex. Neuron. 2019;102(3):668–82 e5. Epub 20190318. doi: 10.1016/j.neuron.2019.02.026. PubMed PMID: 30898376.

75. Page CE, Shepard R, Heslin K, Coutellier L. Prefrontal parvalbumin cells are sensitive to stress and mediate anxiety-related behaviors in female mice. Scientific reports. 2019;9(1):19772. Epub 20191224. doi: 10.1038/s41598-019-56424-9. PubMed PMID: 31875035; PMCID: PMC6930291.

76. Gentet LJ, Kremer Y, Taniguchi H, Huang ZJ, Staiger JF, Petersen CC. Unique functional properties of somatostatin-expressing GABAergic neurons in mouse barrel cortex. Nature neuroscience. 2012;15(4):607–12. Epub 20120226. doi: 10.1038/nn.3051. PubMed PMID: 22366760.

77. Naka A, Adesnik H. Inhibitory Circuits in Cortical Layer 5. Front Neural Circuits. 2016;10:35. Epub 20160506. doi: 10.3389/fncir.2016.00035. PubMed PMID: 27199675; PMCID: PMC4859073.

78. Soumier A, Sibille E. Opposing effects of acute versus chronic blockade of frontal cortex somatostatin-positive inhibitory neurons on behavioral emotionality in mice. Neuropsychopharmacology : official publication of the American College of Neuropsychopharmacology. 2014;39(9):2252–62. Epub 20140401. doi: 10.1038/npp.2014.76. PubMed PMID: 24690741; PMCID: PMC4104344.

79. Abbas AI, Sundiang MJM, Henoch B, Morton MP, Bolkan SS, Park AJ, Harris AZ, Kellendonk C, Gordon JA. Somatostatin Interneurons Facilitate Hippocampal-Prefrontal Synchrony and Prefrontal Spatial Encoding. Neuron. 2018;100(4):926–39 e3. Epub 20181011. doi: 10.1016/j.neuron.2018.09.029. PubMed PMID: 30318409; PMCID: PMC6262834.

80. David C, Schleicher A, Zuschratter W, Staiger JF. The innervation of parvalbumin-containing interneurons by VIP-immunopositive interneurons in the primary somatosensory cortex of the adult rat. The European journal of neuroscience. 2007;25(8):2329–40. doi: 10.1111/j.1460-9568.2007.05496.x. PubMed PMID: 17445231.

81. Pfeffer CK, Xue M, He M, Huang ZJ, Scanziani M. Inhibition of inhibition in visual cortex: the logic of connections between molecularly distinct interneurons. Nature neuroscience. 2013;16(8):1068–76. Epub 20130630. doi: 10.1038/nn.3446. PubMed PMID: 23817549; PMCID: PMC3729586.

82. Pi HJ, Hangya B, Kvitsiani D, Sanders JI, Huang ZJ, Kepecs A. Cortical interneurons that specialize in disinhibitory control. Nature. 2013;503(7477):521-4. Epub 20131006. doi: 10.1038/nature12676. PubMed PMID: 24097352; PMCID: PMC4017628.

83. Lee AT, Cunniff MM, See JZ, Wilke SA, Luongo FJ, Ellwood IT, Ponnavolu S, Sohal VS. VIP Interneurons Contribute to Avoidance Behavior by Regulating Information Flow across Hippocampal-Prefrontal Networks. Neuron. 2019;102(6):1223–34 e4. Epub 20190430. doi: 10.1016/j.neuron.2019.04.001. PubMed PMID: 31053407; PMCID: PMC6800223.

84. Li M, Zhou H, Teng S, Yang G. Activation of VIP interneurons in the prefrontal cortex ameliorates neuropathic pain aversiveness. Cell Rep. 2022;40(11):111333. doi: 10.1016/j.celrep.2022.111333. PubMed PMID: 36103825; PMCID: PMC9520588.

85. Morgan A, Adank D, Johnson K, Butler E, Patel S. 2-Arachidonoylglycerol-mediated endocannabinoid signaling modulates mechanical hypersensitivity associated with alcohol withdrawal in mice. Alcoholism, clinical and experimental research. 2022;46(11):2010–24. Epub 2022/09/21. doi: 10.1111/acer.14949. PubMed PMID: 36125319; PMCID: PMC10091740.

86. Centanni SW, Morris BD, Luchsinger JR, Bedse G, Fetterly TL, Patel S, Winder DG. Endocannabinoid control of the insular-bed nucleus of the stria terminalis circuit regulates negative affective behavior associated with alcohol abstinence. Neuropsychopharmacology : official publication of the American College of Neuropsychopharmacology. 2019;44(3):526–37. Epub 2018/11/06. doi: 10.1038/s41386-018-0257-8. PubMed PMID: 30390064; PMCID: PMC6333805.

87. Marcus DJ, Bedse G, Gaulden AD, Ryan JD, Kondev V, Winters ND, Rosas-Vidal LE, Altemus M, Mackie K, Lee FS, Delpire E, Patel S. Endocannabinoid Signaling Collapse Mediates Stress-Induced Amygdalo-Cortical Strengthening. Neuron. 2020;105(6):1062–76 e6. Epub 2020/01/18. doi: 10.1016/j.neuron.2019.12.024. PubMed PMID: 31948734; PMCID: PMC7992313.

88. Kondev V, Najeed M, Loomba N, Brown J, Winder DG, Grueter BA, Patel S. Synaptic and cellular endocannabinoid signaling mechanisms regulate stress-induced plasticity of nucleus accumbens somatostatin neurons. Proceedings of the National Academy of Sciences of the United States of America. 2023;120(34):e2300585120. Epub 2023/08/17. doi: 10.1073/pnas.2300585120. PubMed PMID: 37590414; PMCID: PMC10450650.

89. Ting JT, Lee BR, Chong P, Soler-Llavina G, Cobbs C, Koch C, Zeng H, Lein E. Preparation of Acute Brain Slices Using an Optimized N-Methyl-D-glucamine Protective Recovery Method. J Vis Exp. 2018(132). Epub 20180226. doi: 10.3791/53825. PubMed PMID: 29553547; PMCID: PMC5931343.

90. Guzman SJ, Schlogl A, Schmidt-Hieber C. Stimfit: quantifying electrophysiological data with Python. Front Neuroinform. 2014;8:16. Epub 2014/03/07. doi: 10.3389/fninf.2014.00016. PubMed PMID: 24600389; PMCID: PMC3931263.

91. Pronneke A, Scheuer B, Wagener RJ, Mock M, Witte M, Staiger JF. Characterizing VIP Neurons in the Barrel Cortex of VIPcre/tdTomato Mice Reveals Layer-Specific Differences. Cereb Cortex. 2015;25(12):4854–68. Epub 20150924. doi: 10.1093/cercor/bhv202. PubMed PMID: 26420784; PMCID: PMC4635925.

92. Rudy B, Fishell G, Lee S, Hjerling-Leffler J. Three groups of interneurons account for nearly 100% of neocortical GABAergic neurons. Dev Neurobiol. 2011;71(1):45–61. doi: 10.1002/dneu.20853. PubMed PMID: 21154909; PMCID: PMC3556905.

93. Minnig MA, Park T, Echeveste Sanchez M, Cottone P, Sabino V. Viral-Mediated Knockdown of Nucleus Accumbens Shell PAC1 Receptor Promotes Excessive Alcohol Drinking in Alcohol-Preferring Rats. Front Behav Neurosci. 2021;15:787362. Epub 20211203. doi: 10.3389/fnbeh.2021.787362. PubMed PMID: 34924973; PMCID: PMC8678417.

94. Torres Irizarry VC, Feng B, Yang X, Patel N, Schaul S, Ibrahimi L, Ye H, Luo P, Carrillo-Saenz L, Lai P, Kota M, Dixit D, Wang C, Lasek AW, He Y, Xu P. Estrogen signaling in the dorsal raphe regulates binge-like drinking in mice. Transl Psychiatry. 2024;14(1):122. Epub 20240227. doi: 10.1038/s41398-024-02821-2. PubMed PMID: 38413577; PMCID: PMC10899193.

95. Deal AL, Bass CE, Grinevich VP, Delbono O, Bonin KD, Weiner JL, Budygin EA. Bidirectional Control of Alcohol-drinking Behaviors Through Locus Coeruleus Optoactivation. Neuroscience. 2020;443:84–92. Epub 20200721. doi: 10.1016/j.neuroscience.2020.07.024. PubMed PMID: 32707291; PMCID: PMC8074022.

96. Faccidomo S, Bannai M, Miczek KA. Escalated aggression after alcohol drinking in male mice: dorsal raphe and prefrontal cortex serotonin and 5-HT(1B) receptors. Neuropsychopharmacology : official publication of the American College of Neuropsychopharmacology. 2008;33(12):2888–99. Epub 20080227. doi: 10.1038/npp.2008.7. PubMed PMID: 18305458.

97. Packer AM, Yuste R. Dense, unspecific connectivity of neocortical parvalbumin-positive interneurons: a canonical microcircuit for inhibition? The Journal of neuroscience : the official journal of the Society for Neuroscience. 2011;31(37):13260–71. doi: 10.1523/JNEUROSCI.3131-11.2011. PubMed PMID: 21917809; PMCID: PMC3178964.

98. Isaacson JS, Scanziani M. How inhibition shapes cortical activity. Neuron. 2011;72(2):231–43. doi: 10.1016/j.neuron.2011.09.027. PubMed PMID: 22017986; PMCID: PMC3236361.

99. Thompson SM, Ferranti AS, Joffe ME. Acute alcohol and chronic drinking bidirectionally regulate the excitability of prefrontal cortex vasoactive intestinal peptide interneurons. bioRxiv. 2023. Epub 2023/03/23. doi: 10.1101/2023.03.07.531614. PubMed PMID: 36945582; PMCID: PMC10028880 R00AA027806], the Whitehall Foundation [grant number 2022-08-005], and the Brain and Behavior Research Foundation. The authors declare no potential conflicts of interest.

100. Wu J, Zhao Z, Shi Y, He M. Cortical VIP(+) Interneurons in the Upper and Deeper Layers Are Transcriptionally Distinct. J Mol Neurosci. 2022;72(8):1779–95. Epub 20220616. doi: 10.1007/s12031-022-02040-8. PubMed PMID: 35708842.

101. Gouwens NW, Sorensen SA, Baftizadeh F, Budzillo A, Lee BR, Jarsky T, Alfiler L, Baker K, Barkan E, Berry K, Bertagnolli D, Bickley K, Bomben J, Braun T, Brouner K, Casper T, Crichton K, Daigle TL, Dalley R, de Frates RA, Dee N, Desta T, Lee SD, Dotson N, Egdorf T, Ellingwood L, Enstrom R, Esposito L, Farrell C, Feng D, Fong O, Gala R, Gamlin C, Gary A, Glandon A, Goldy J, Gorham M, Graybuck L, Gu H, Hadley K, Hawrylycz MJ, Henry AM, Hill D, Hupp M, Kebede S, Kim TK, Kim L, Kroll M, Lee C, Link KE, Mallory M, Mann R, Maxwell M, McGraw M, McMillen D, Mukora A, Ng L, Ng L, Ngo K, Nicovich PR, Oldre A, Park D, Peng H, Penn O, Pham T, Pom A, Popovic Z, Potekhina L, Rajanbabu R, Ransford S, Reid D, Rimorin C, Robertson M, Ronellenfitch K, Ruiz A, Sandman D, Smith K, Sulc J, Sunkin SM, Szafer A, Tieu M, Torkelson A, Trinh J, Tung H, Wakeman W, Ward K, Williams G, Zhou Z, Ting JT, Arkhipov A, Sumbul U, Lein ES, Koch C, Yao Z, Tasic B, Berg J, Murphy GJ, Zeng H. Integrated Morphoelectric and Transcriptomic Classification of Cortical GABAergic Cells. Cell. 2020;183(4):935–53 e19. doi: 10.1016/j.cell.2020.09.057. PubMed PMID: 33186530; PMCID: PMC7781065.

102. Egli M, Koob GF, Edwards S. Alcohol dependence as a chronic pain disorder. Neuroscience and biobehavioral reviews. 2012;36(10):2179–92. Epub 2012/09/15. doi: 10.1016/j.neubiorev.2012.07.010. PubMed PMID: 22975446; PMCID: 3612891.

103. Koob GF. Addiction is a Reward Deficit and Stress Surfeit Disorder. Frontiers in psychiatry. 2013;4:72. Epub 2013/08/06. doi: 10.3389/fpsyt.2013.00072. PubMed PMID: 23914176; PMCID: 3730086.

104. Ruelas M, Medina-Ceja L, Fuentes-Aguilar RQ. A scoping review of the relationship between alcohol, memory consolidation and ripple activity: An overview of common methodologies to analyse ripples. The European journal of neuroscience. 2023;58(10):4137–54. Epub 20231012. doi: 10.1111/ejn.16168. PubMed PMID: 37827165.

105. Robins MT, Heinricher MM, Ryabinin AE. From Pleasure to Pain, and Back Again: The Intricate Relationship Between Alcohol and Nociception. Alcohol Alcohol. 2019;54(6):625–38. Epub 2019/09/12. doi: 10.1093/alcalc/agz067. PubMed PMID: 31509854.

106. Centanni SW, Bedse G, Patel S, Winder DG. Driving the Downward Spiral: Alcohol-Induced Dysregulation of Extended Amygdala Circuits and Negative Affect. Alcoholism, clinical and experimental research. 2019;43(10):2000–13. Epub 20190830. doi: 10.1111/acer.14178. PubMed PMID: 31403699; PMCID: PMC6779502.

107. Jones AF, Sheets PL. Sex-Specific Disruption of Distinct mPFC Inhibitory Neurons in Spared-Nerve Injury Model of Neuropathic Pain. Cell Rep. 2020;31(10):107729. doi: 10.1016/j.celrep.2020.107729. PubMed PMID: 32521254; PMCID: PMC7372908.

108. Zhang Z, Gadotti VM, Chen L, Souza IA, Stemkowski PL, Zamponi GW. Role of Prelimbic GABAergic Circuits in Sensory and Emotional Aspects of Neuropathic Pain. Cell Rep. 2015;12(5):752–9. Epub 20150723. doi: 10.1016/j.celrep.2015.07.001. PubMed PMID: 26212331.

109. Nawreen N, Oshima K, Chambers J, Smail M, Herman JP. Inhibition of prefrontal cortex parvalbumin interneurons mitigates behavioral and physiological sequelae of chronic stress in male mice. Stress. 2024;27(1):2361238. Epub 20240704. doi: 10.1080/10253890.2024.2361238. PubMed PMID: 38962839.

110. Fogaca MV, Wu M, Li C, Li XY, Picciotto MR, Duman RS. Inhibition of GABA interneurons in the mPFC is sufficient and necessary for rapid antidepressant responses. Molecular psychiatry. 2021;26(7):3277–91. Epub 20201017. doi: 10.1038/s41380-020-00916-y. PubMed PMID: 33070149; PMCID: PMC8052382.

111. Hatter JA, Scott MM. Selective ablation of VIP interneurons in the rodent prefrontal cortex results in increased impulsivity. PloS one. 2023;18(6):e0286209. Epub 20230602. doi: 10.1371/journal.pone.0286209. PubMed PMID: 37267385; PMCID: PMC10237669.

112. Medina KL, McQueeny T, Nagel BJ, Hanson KL, Schweinsburg AD, Tapert SF. Prefrontal cortex volumes in adolescents with alcohol use disorders: unique gender effects. Alcoholism, clinical and experimental research. 2008;32(3):386–94. doi: 10.1111/j.1530-0277.2007.00602.x. PubMed PMID: 18302722; PMCID: PMC2825148.

113. Squeglia LM, Sorg SF, Schweinsburg AD, Wetherill RR, Pulido C, Tapert SF. Binge drinking differentially affects adolescent male and female brain morphometry. Psychopharmacology. 2012;220(3):529–39. Epub 20110928. doi: 10.1007/s00213-011-2500-4. PubMed PMID: 21952669; PMCID: PMC3527131.

114. Seo S, Beck A, Matthis C, Genauck A, Banaschewski T, Bokde ALW, Bromberg U, Buchel C, Quinlan EB, Flor H, Frouin V, Garavan H, Gowland P, Ittermann B, Martinot JL, Paillere Martinot ML, Nees F, Papadopoulos Orfanos D, Poustka L, Hohmann S, Frohner JH, Smolka MN, Walter H, Whelan R, Desrivieres S, Heinz A, Schumann G, Obermayer K. Risk profiles for heavy drinking in adolescence: differential effects of gender. Addiction biology. 2019;24(4):787–801. Epub 20180530. doi: 10.1111/adb.12636. PubMed PMID: 29847018.

115. Squeglia LM, Schweinsburg AD, Pulido C, Tapert SF. Adolescent binge drinking linked to abnormal spatial working memory brain activation: differential gender effects. Alcoholism, clinical and experimental research. 2011;35(10):1831–41. Epub 20110718. doi: 10.1111/j.1530-0277.2011.01527.x. PubMed PMID: 21762178; PMCID: PMC3183294.

116. Acker C. Neuropsychological deficits in alcoholics: the relative contributions of gender and drinking history. Br J Addict. 1986;81(3):395–403. doi: 10.1111/j.1360-0443.1986.tb00346.x. PubMed PMID: 3461848.

117. Niaura RS, Nathan PE, Frankenstein W, Shapiro AP, Brick J. Gender differences in acute psychomotor, cognitive, and pharmacokinetic response to alcohol. Addictive behaviors. 1987;12(4):345–56. doi: 10.1016/0306-4603(87)90048-7. PubMed PMID: 3687517.

118. Nixon SJ. Cognitive Deficits in Alcoholic Women. Alcohol Health Res World. 1994;18(3):228–32. PubMed PMID: 31798100; PMCID: PMC6876400.

119. Flannery B, Fishbein D, Krupitsky E, Langevin D, Verbitskaya E, Bland C, Bolla K, Egorova V, Bushara N, Tsoy M, Zvartau E. Gender differences in neurocognitive functioning among alcohol-dependent Russian patients. Alcoholism, clinical and experimental research. 2007;31(5):745–54. Epub 20070326. doi: 10.1111/j.1530-0277.2007.00372.x. PubMed PMID: 17386068.

120. Scaife JC, Duka T. Behavioural measures of frontal lobe function in a population of young social drinkers with binge drinking pattern. Pharmacology, biochemistry, and behavior. 2009;93(3):354–62. Epub 20090602. doi: 10.1016/j.pbb.2009.05.015. PubMed PMID: 19497334.

121. Kirsch DE, Belnap MA, Burnette EM, Grodin EN, Ray LA. Pharmacological Treatments for Alcohol Use Disorder: Considering the Role of Sex and Gender. Current Addiction Reports. 2024;11(1):81–93. doi: 10.1007/s40429-023-00535-x. PubMed PMID: WOS:001132685700001.

122. Zindel LR, Kranzler HR. Pharmacotherapy of alcohol use disorders: seventy-five years of progress. J Stud Alcohol Drugs Suppl. 2014;75(17):79–88. doi: 10.15288/jsads.2014.s17.79. PubMed PMID: 24565314; PMCID: PMC4453501.

123. Hasin DS, Stinson FS, Ogburn E, Grant BF. Prevalence, correlates, disability, and comorbidity of DSM-IV alcohol abuse and dependence in the United States: results from the National Epidemiologic Survey on Alcohol and Related Conditions. Archives of general psychiatry. 2007;64(7):830–42. Epub 2007/07/04. doi: 10.1001/archpsyc.64.7.830. PubMed PMID: 17606817.

