## Supplemental Figures for "Alcohol Withdrawal Alters the Inhibitory Landscape of the Prelimbic Cortex in an Interneuron- and Sex-specific Manner"

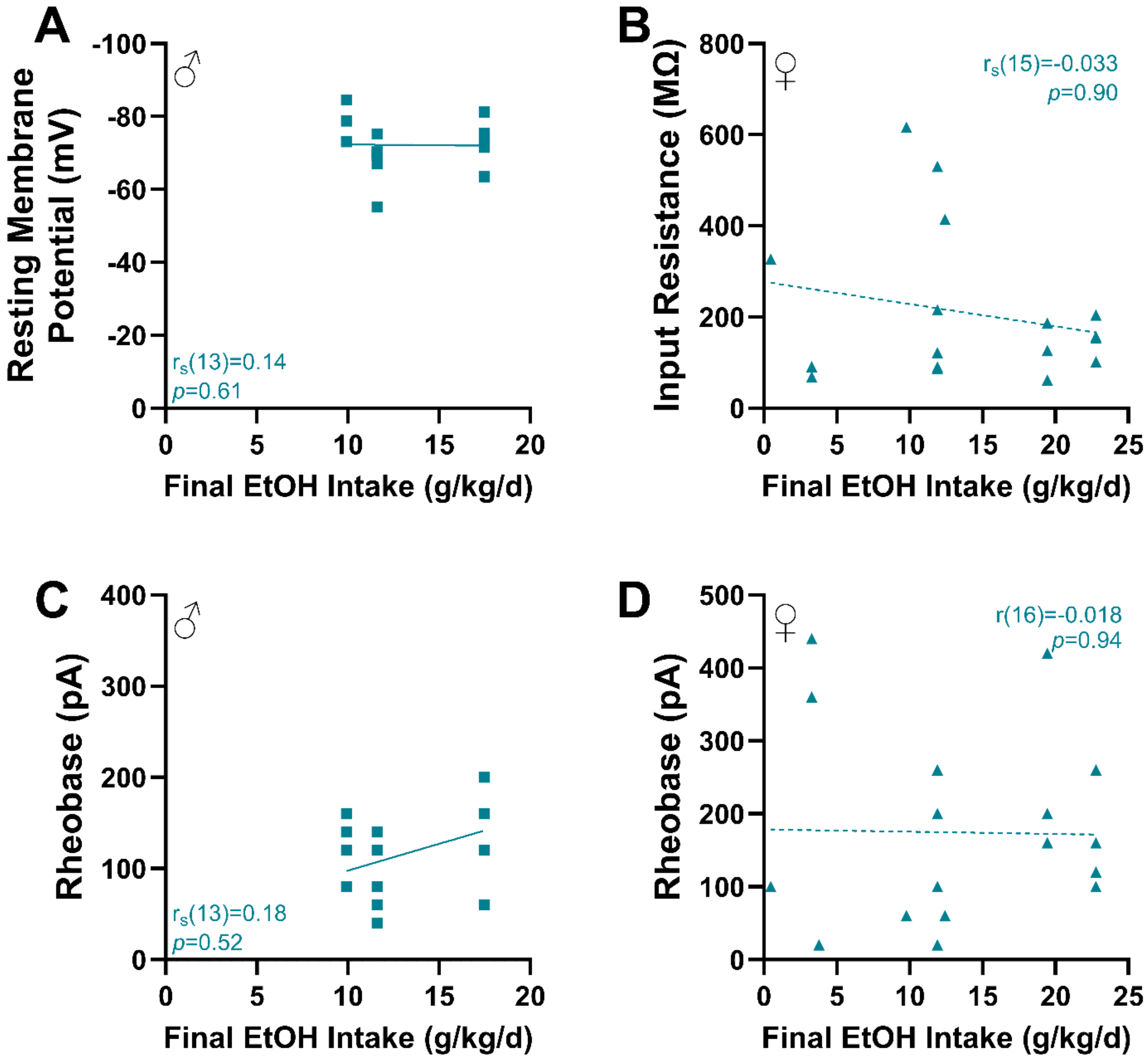


<u>Figure S1:</u> **Non-significant correlations between EtOH intake and neuronal excitability of prelimbic parvalbumin (PV) interneurons. A:** Resting membrane potential (males). **B:** Input resistance (females). **C:** Rheobase (males). **D:** Rheobase (females). Data are shown as individual points with no significant correlations observed.

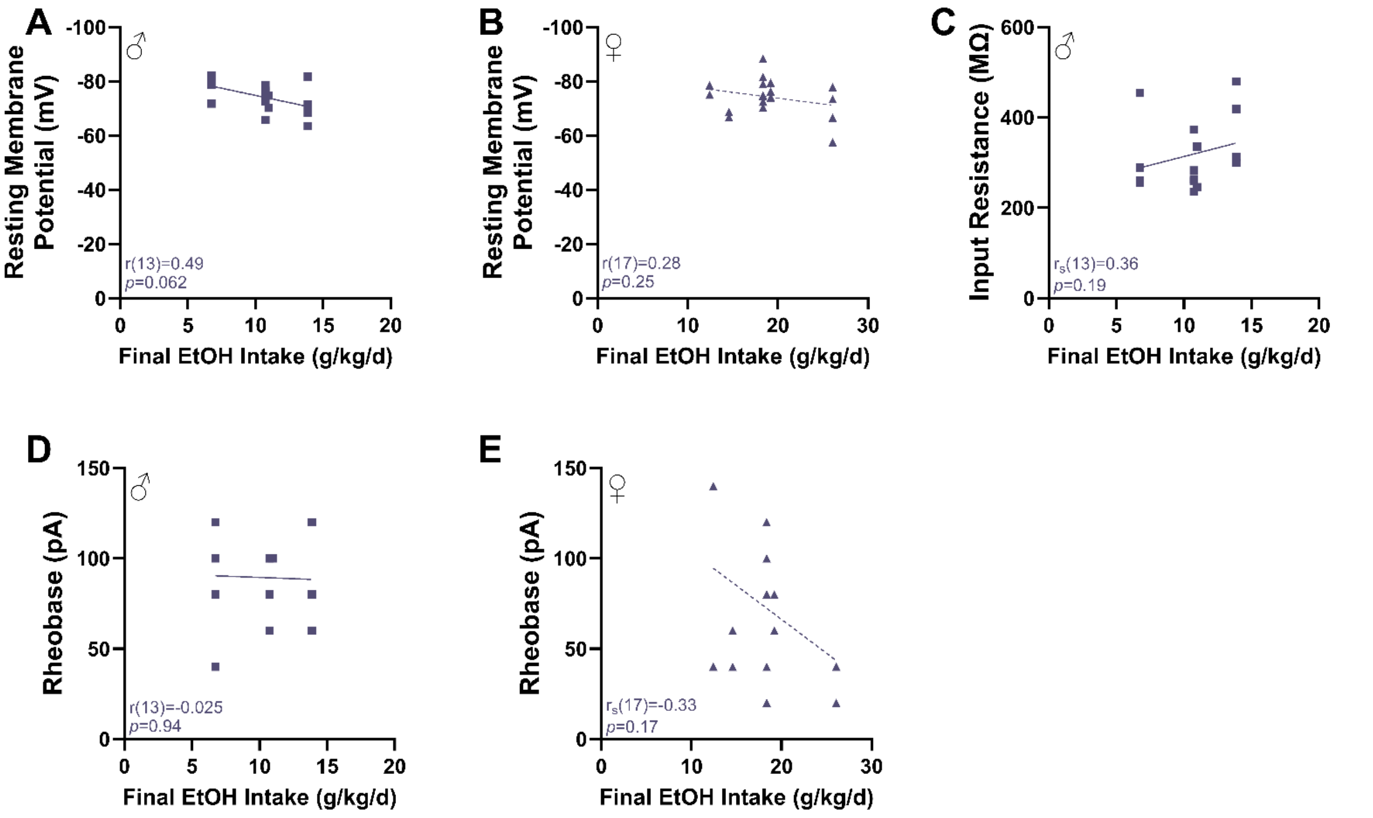


<u>Figure S2:</u> **Non-significant correlations between EtOH intake and neuronal excitability of prelimbic somatostatin (SOM) interneurons. A:** Resting membrane potential (males). **B:** Resting membrane potential (females). **C:** Input resistance (males). **D:** Rheobase (males). **E:** Rheobase (females). Data are shown as individual points with no significant correlations observed.

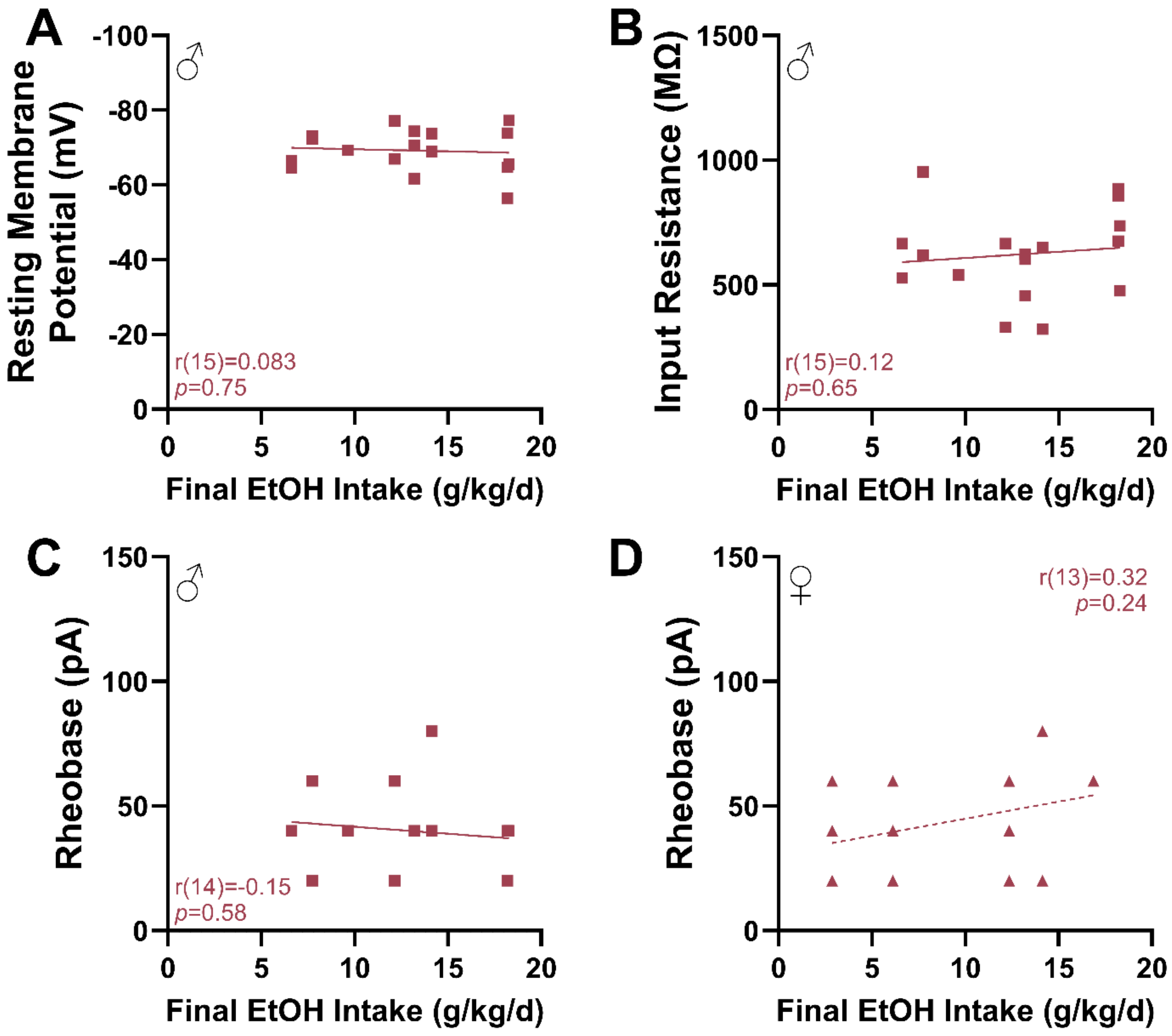


**<u>Figure S3:</u> Non-significant correlations between EtOH intake and neuronal excitability of prelimbic vasoactive intestinal peptide (VIP) interneurons. A:** Resting membrane potential (males). **B**: Input resistance (males). **C:** Rheobase (males). **D:** Rheobase (females). Data are shown as individual points with no significant correlations observed.

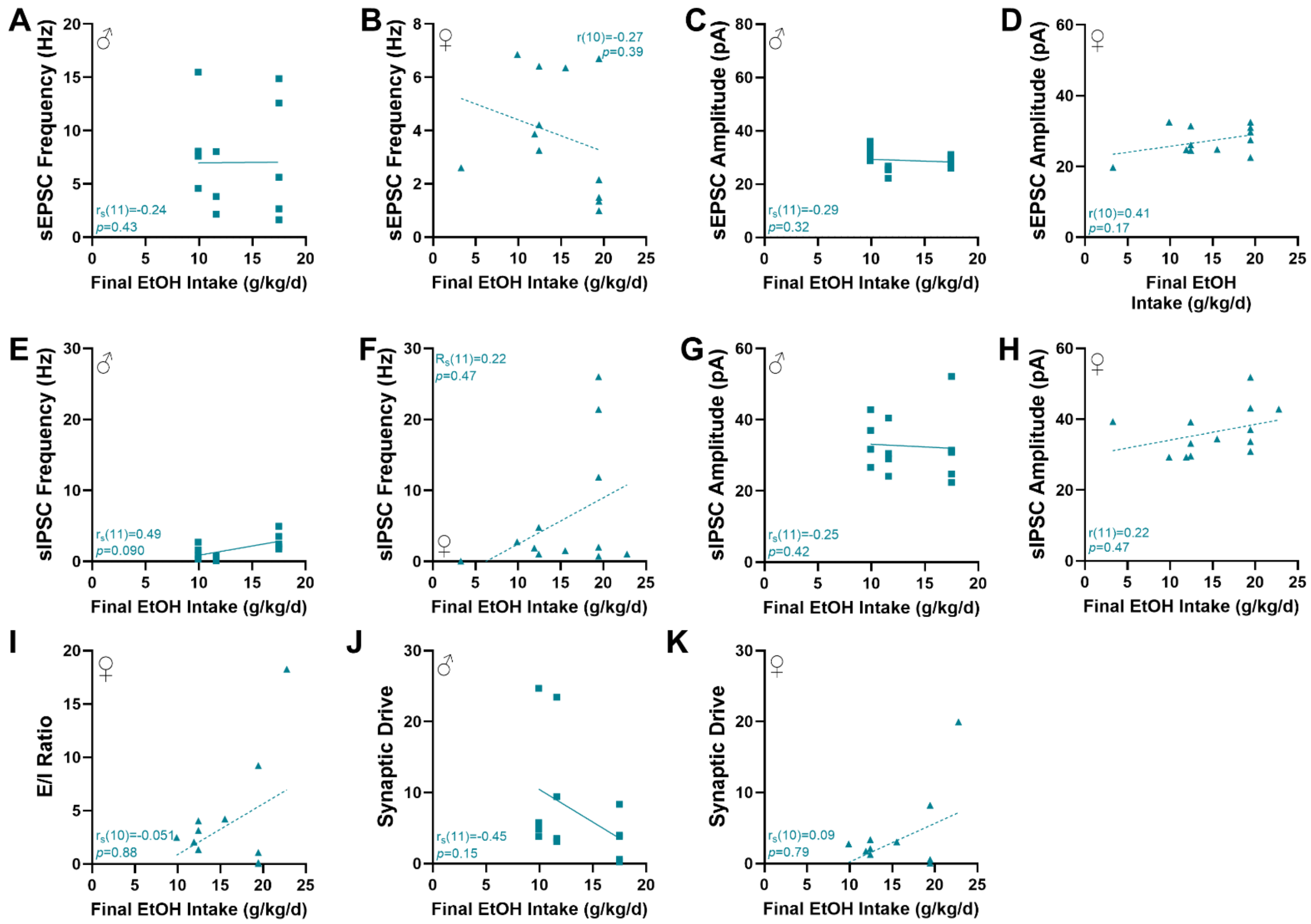


**<u>Figure S4:</u> Non-significant correlations between EtOH intake and synaptic transmission in prelimbic parvalbumin (PV) neurons. A:** sEPSC frequency (males). **B:** sEPSC frequency (females). **C:** sEPSC amplitude (males). **D:** sEPSC amplitude (females). **E:** sIPSC frequency (males). **F:** sIPSC frequency (females). **G:** sIPSC amplitude (males). **H:** sIPSC amplitude (females). **I:** E/I Ratio (females). **J:** Synaptic drive (males). **K:** Synaptic drive (females). Data are shown as individual points with no significant correlations observed.

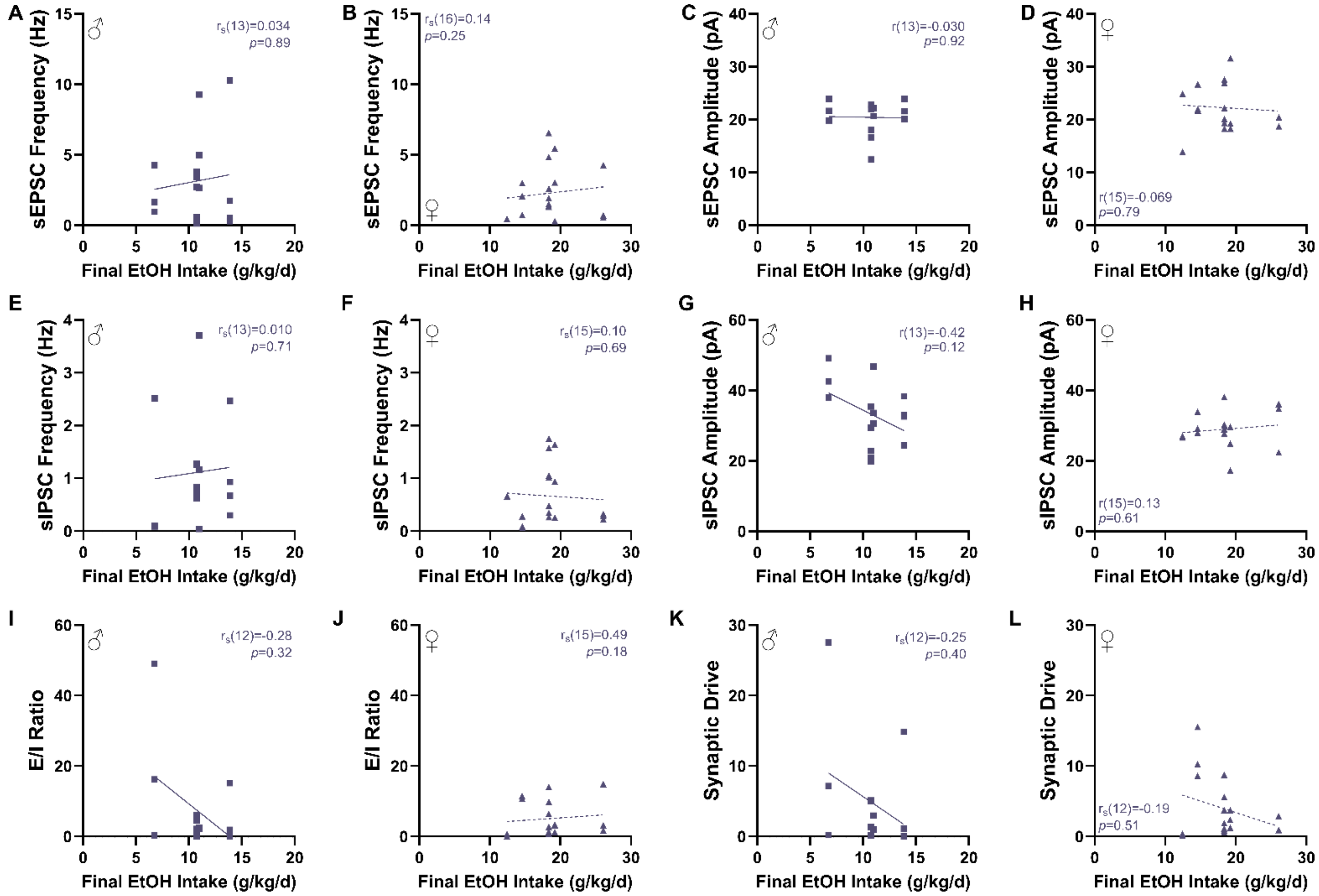


**<u>Figure S5:</u> Non-significant correlations between EtOH intake and synaptic transmission in prelimbic somatostatin (SOM) neurons. A:** sEPSC frequency (males). **B:** sEPSC frequency (females). **C:** sEPSC amplitude (males). **D:** sEPSC amplitude (females). **E:** sIPSC frequency (males). **F:** sIPSC frequency (females). **G:** sIPSC amplitude (males). **H:** sIPSC amplitude (females). **I:** E/I Ratio (males). **J:** E/I Ratio (females). **K:** Synaptic drive (males). **L:** Synaptic drive (females). Data are shown as individual points with no significant correlations observed.

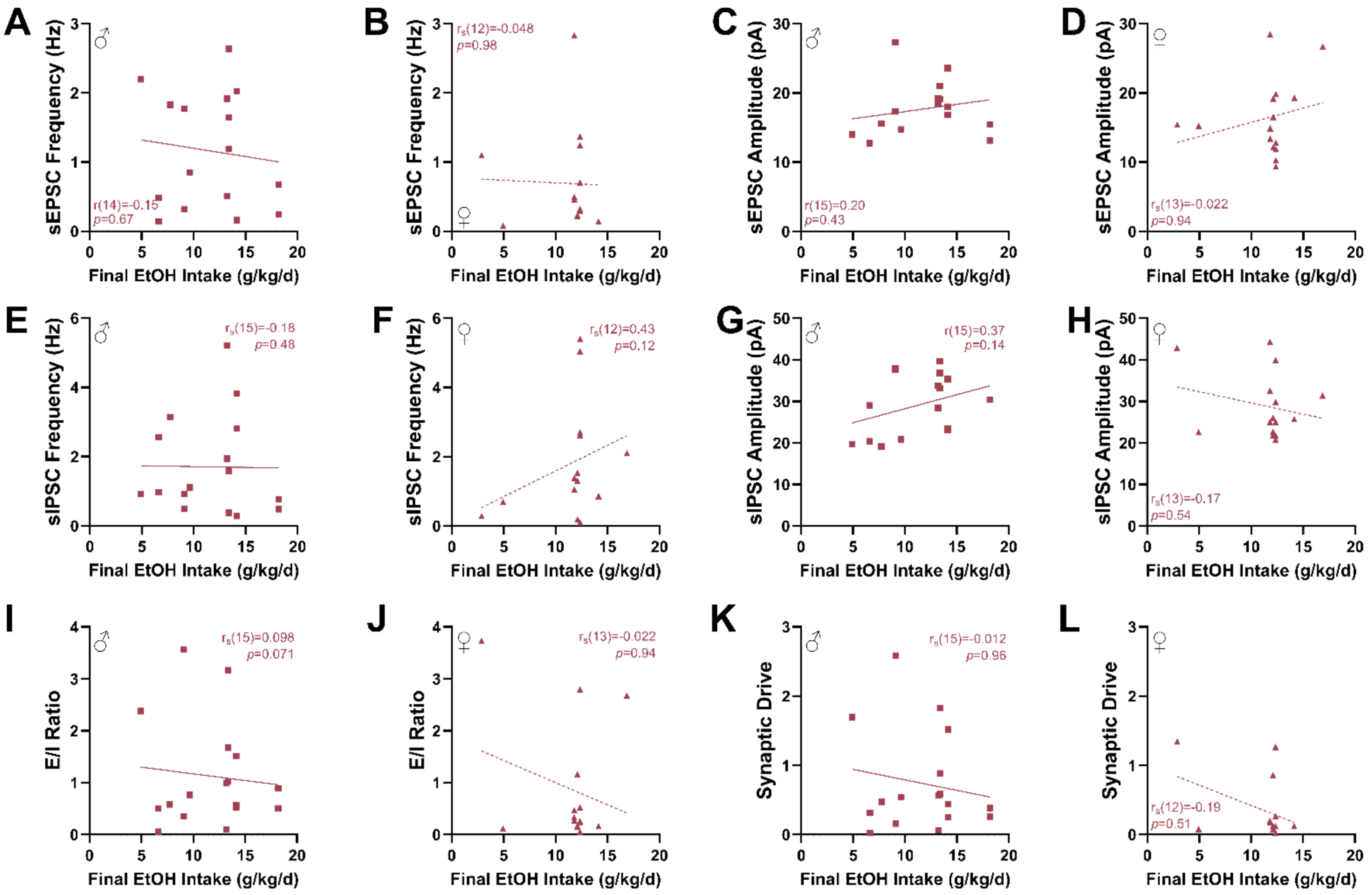


**Figure S6: Non-significant correlations between EtOH intake and synaptic transmission in prelimbic vasoactive intestinal peptide (VIP) neurons. A:** sEPSC frequency (males). **B:** sEPSC frequency (females). **C:** sEPSC amplitude (males). **D:** sEPSC amplitude (females). **E:** sIPSC frequency (males). **F:** sIPSC frequency (females). **G:** sIPSC amplitude (males). **H:** sIPSC (females). **I:** E/I Ratio (males). **J:** E/I Ratio (females). **K:** Synaptic drive (males). **L:** Synaptic drive (females). Data are shown as individual points with no significant correlations observed.
